# Design and characterization of a protein fold switching network

**DOI:** 10.1101/2022.10.26.513944

**Authors:** Biao Ruan, Yanan He, Yingwei Chen, Eun Jung Choi, Yihong Chen, Dana Motabar, Tsega Solomon, Richard Simmerman, Thomas Kauffman, D. Travis Gallagher, John Orban, Philip N. Bryan

**Author notes:** B.R., Y.H., and Yw.C. contributed equally. To whom correspondence should be addressed: Philip N. Bryan, John Orban, **Email:**.

## Abstract

Protein sequences encoding three common small folds (3α, β–grasp, and α/β–plait) were connected in a network with high-identity intersections, termed nodes. The structures of proteins around nodes were determined using NMR spectroscopy and analyzed for stability and binding function. To generate nodes, the amino acid sequence encoding a shorter fold (3a or β–grasp) is embedded in the structure of the ^~^50% longer α/β–plait fold and a new sequence is designed that satisfies two sets of native interactions. This leads to protein pairs with a 3a or β–grasp fold in the shorter form but an α/β–plait fold in the longer form. Further, embedding smaller antagonistic folds in longer folds creates critical states in the longer folds such that single amino acid substitutions can switch both their fold and function. This suggests that abrupt fold switching may be a mechanism of evolving new protein structures and functions.

## INTRODUCTION

Fold switching occurs when one amino acid sequence has a propensity for two completely different, but well-ordered, conformations. Many examples of both natural and engineered fold switching demonstrate that proteins can have a stable native fold while simultaneously hiding latent propensities for alternative states with new functions ^1–8^. This fact has many implications for understanding how amino acid sequence encodes structure, how proteins evolve, how mutation is related to disease, and how function is annotated to sequences of unknown structure ^9–22^.

In this work, a network of high-identity intersections (nodes) was engineered that connects three common and well-studied protein folds. Two of these folds are from Streptococcal Protein G which contains two types of domains that bind to serum proteins in blood: the G_A_ domain binds to human serum albumin (HSA) ^23,24^ and the G_B_ domain binds to the constant (Fc) region of IgG ^25,26^. The third protein is S6, a component of the 30S ribosomal subunit of *Thermus thermophilus* ^27–31^. For simplicity, the S6 fold is referred to as an S-fold, the G_A_ fold as an A-fold, and the G_B_ fold as a B-fold. These proteins share no significant homology and are representative of three of the ten most common folds: the S-fold is a thioredoxin-like α/β plait; the A-fold is a homeodomain-like 3α–helix bundle; the B-fold is a ubiquitin-like β grasp^32^.

**Fig. 1** depicts a map of the nodes that were engineered here. The arrows in **Fig. 1** show a network originating with the natural S6 sequence. Circles represent nodes in the network at which structural and/or functional switches occur. The SI and S’I nodes are a branch point and lead down diverging sequence pathways, one leading to a node with the A-fold (S/A) and one to a node with the B-fold (S/B). Intersecting mutational pathways lead from S/A to the native G_A_ protein and S/B to the native G_B_ protein. At this intersection (A/B), an A-fold switches to a B-fold.

**Fig. 1:**
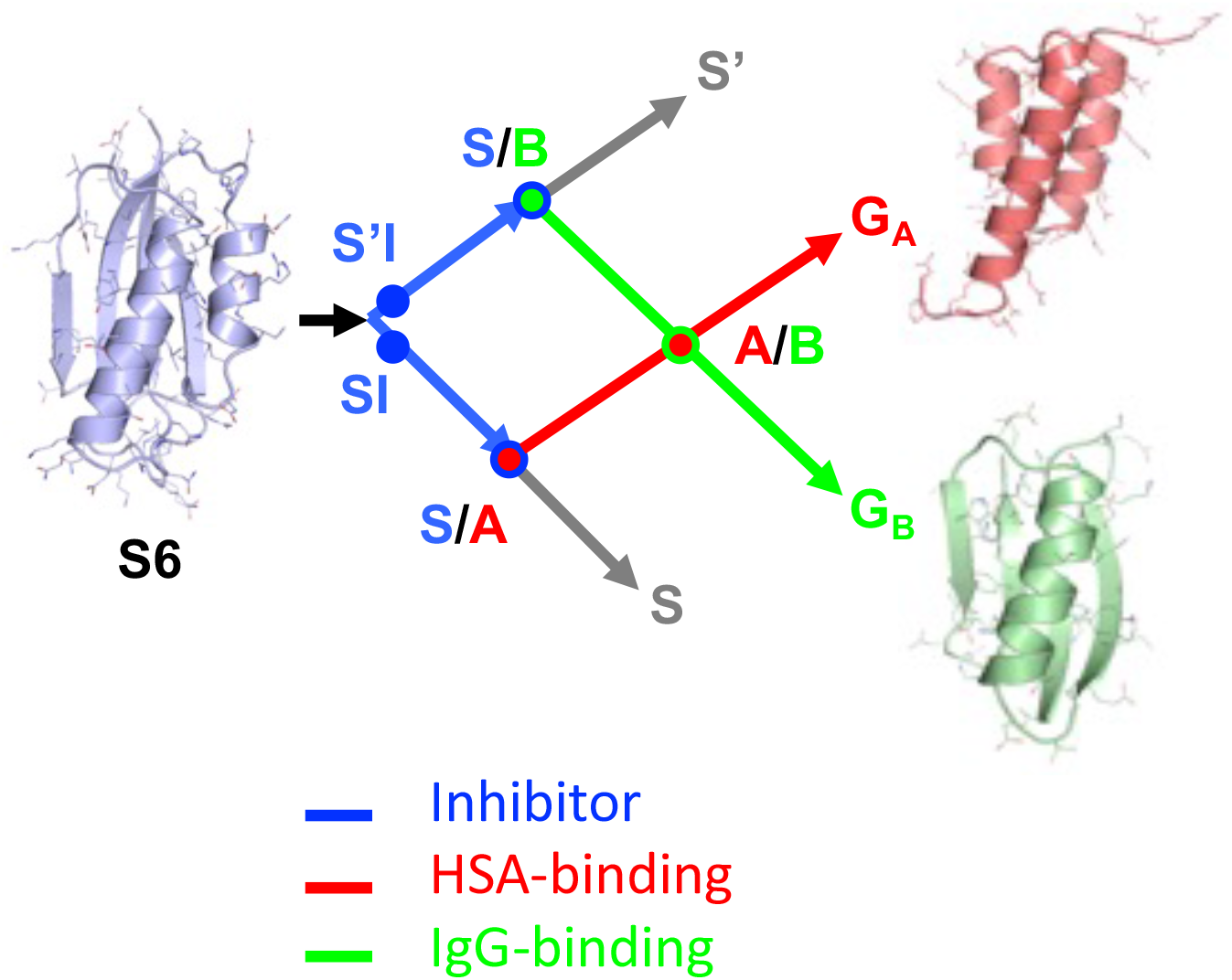
Overview of engineered nodes in the S6, G_A_, and G_B_ network. S6 is the origin sequence in the engineering process. SI and S’I are separate nodes and are loop size variants of the S-fold, both having protease inhibitor function. The SI branch of the mutational path leads to a node with the A-fold and HSA binding function. The S’I branch of the path leads to a node with the B-fold and IgG binding function. The S/A node (blue and red circle) includes proteins S_a1_, S_a2_, A_1_, and A_2_. The S/B node (blue and green circle) includes proteins S_b3_, S_b4_, S_b5_, B_3_, and B_4_. The A and B paths themselves intersect at an A/B node (green and red circle) at which A- and B-folds are nearly iso-energetic and bifunctional. The S and S’ branches continue and connect with many other natural sequences in the α/β plait super-fold family.

Proteins around the A/B node have been extensively characterized in our earlier work and demonstrated protein structure can be encoded by a small number of essential residues, and that a very limited subset of intra-protein interactions can tip the balance from one fold and function to another ^33–35^. Here we determine that both G_A_ and G_B_ can switch into a third fold (α/β–plait) and show that three folds and four functions can be connected in a network that avoids unfolded and functionless states. We describe how these nodes were engineered, determine key structures using NMR spectroscopy, and analyze stability and binding function. The ability to design and characterize nodes connecting three common small folds suggests that fold switching may be a general phenomenon in the evolution of protein structure and function.

## RESULTS

### Designing a functional switch from ribosomal protein to protease inhibitor

The S6 ribosomal protein is structurally homologous to subtilisin protease inhibitors known as prodomains (**Fig. 2A, B**) ^36^. Prodomain-type inhibitors have two binding surfaces with the protease. One surface comprises the last nine C-terminal amino acids of the inhibitor which bind in the substrate binding cleft of the protease (**Fig. 2B**). A second, more dynamic surface is formed between two subtilisin helices and the large surface of the β–sheet in the α/β-plait topology of the inhibitor (**Fig. 2B**) ^37–39^. As a result, the S6 protein could be converted into a subtilisin inhibitor protein of the same overall fold (denoted SI) by replacing its nine C-terminal amino acids with residues optimized to bind in the substrate binding cleft of subtilisin. This replacement results in new contacts between the SI β–sheet and the subtilisin surface helices (**Fig. 2B**).

**Fig. 2:**
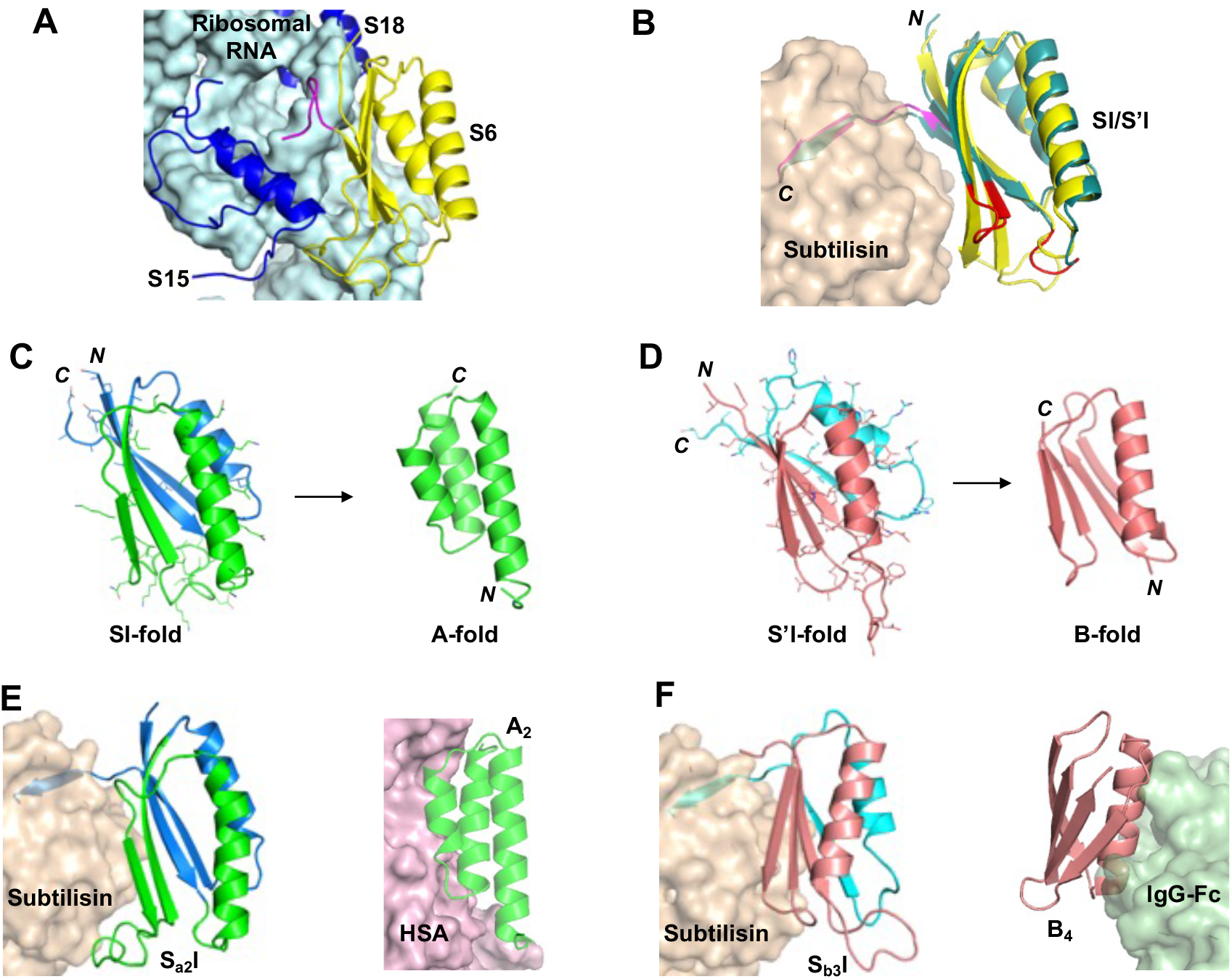
Summary of switches in structure and function. A) Structure of the S6 protein (yellow), RNA (light blue), and S15 and S18 proteins (blue) in the 30S ribosome (PDB 1FKA). C-terminal amino acids of S6 are in magenta. B) Subtilisin (wheat) is shown in complex with a model of the SI-inhibitor (yellow). The C-terminal nine amino acids of SI are shown in magenta. These positions were mutated in native S6 to generate affinity for subtilisin. The S’I-inhibitor (teal) is also shown with the altered loops in red. The subtilisin used in the modeling was the engineered RAS-specific protease. C) The S_a1_ protein (blue and green) was generated from SI by mutating the 45 positions (mutant side chains shown with sticks). Deletion of C-terminal amino acids (blue) switches S_a1_ into the A-fold (green). D) The S_b3_ protein (rose and cyan) was generated from S’I by mutating the 67 positions (mutant side chains shown with sticks). Deletion of C-terminal amino acids (cyan) or point mutation will switch S_b3_ into a B-fold (rose). E) Model of S_a2_I (green and blue) bound to subtilisin (wheat). Model of A_2_ (based on A_1_ structure) bound to HSA (violet). The HSA complex used PDB 2VDB as the template. F) Model of S_b3_ (rose and cyan) bound to subtilisin (wheat). Model of B_4_ (rose) bound to Fc (mint). The Fc complex used PDB 1FCC as the template. The subtilisin used in the modeling and inhibition measurements was the engineered RAS-specific protease (PDB 6UAO).

The SI-protein is 99 amino acids in length and has a 10 residue loop between β2 and β3. However, there are many natural variations in the length of loops in the conserved α/β-plait topology ^40^. Therefore, we also engineered a 91 amino acid version of the S-fold (denoted S’I), which resembles the topology of natural prodomain inhibitors (**Fig. S1**). Specifically, the S’I inhibitor has a longer loop connecting β1 to α1 and a shorter turn connecting β2 to β3 (**Fig. 2B**).

The SI and S’I proteins were expressed and purified by binding to a protease column ^41^. The CD spectra were compared to the native S6 protein (**Fig. S1**). Inhibition constants (K_I_) were measured using an engineered RAS-specific subtilisin protease and the peptide substrate QEEYSAM-AMC ^41^. SI and S’I inhibit the RAS-specific protease with K_I_ values of 200nM and 60nM, respectively (**Table S1**). The details of the competitive inhibition assay are described in Materials and Methods. The results demonstrate that a ribosomal protein can be converted into a protease inhibitor with minor modification (and without a fold switch). In addition, however, the SI and S’I proteins also facilitated engineering subsequent switches to new folds and functions by linking each of the S-, A-, and B- folds to easily measured binding functions: protease inhibition (S or S’-fold); HSA-binding (A-fold); and IgG binding (B-fold).

### Designing fold switches

In previous work we created sequences that populate both A- and B-folds by threading the A-sequence through the B-fold, finding a promising alignment, and then using phage-display selection to reconcile one sequence to both folds ^34,42,43^. Here the approach is conceptually similar, except that we use Rosetta ^44^ as a computational design tool to test compatible mutations rather than phage display. The design process is as follows:

i. Thread the A- or B- sequence through both SI and S’I-fold types.
ii. Identify alignments that minimize the number of catastrophic interactions.
iii. Design mutations to resolve unfavorable interactions in clusters of 4-6 amino acids using Pymol ^45^ and energy minimize using Rosetta-Relax ^44^.
iv. Optimize protein stability in the S-fold by computationally mutating amino acids at non-overlapping positions. Repeat energy minimization and evaluation with Rosetta-Relax.
v. To reduce uncertainties involved in computational design, conserve original amino acids whenever possible.

There is no reason to assume that this method is optimal. We are just applying a practicable scheme for engineering sequences compatible with two sets of native interactions and then evaluating structure, stability, and function. Initial designs were refined based on structural analysis with NMR, thermodynamic analysis of unfolding, and functional analysis using binding assays, as described below. All designed proteins were expressed in *E. coli* and purified to homogeneity as described in Materials and Methods.

### Designing a switch from α/β-plait protease inhibitor to 3α HSA-binding protein

Alignment of the 56 amino acid HSA-binding, A-fold with the 99 amino acid SI-fold and subsequent mutation to resolve catastrophic interactions produced low energy switch candidates denoted S_a1_ and A_1_. The exact sequence of A_1_ is embedded in S_a1_ at positions 11-66 such that the α1 helices are structurally aligned (**Fig. 3A**, **Fig. S2A**). Their final computational models were generated by Rosetta using the Relax application. The Relax protocol searches the local conformational space around an experimentally-determined structure and is used only to evaluate whether the designed mutations have favorable native interactions within that limited conformational space. The designed models of S_a1_ and A_1_ are very similar in energy compared to the respective relaxed native structures (**Fig. S3 and Supplemental PDB files of the Rosetta models**).

**Fig. 3:**
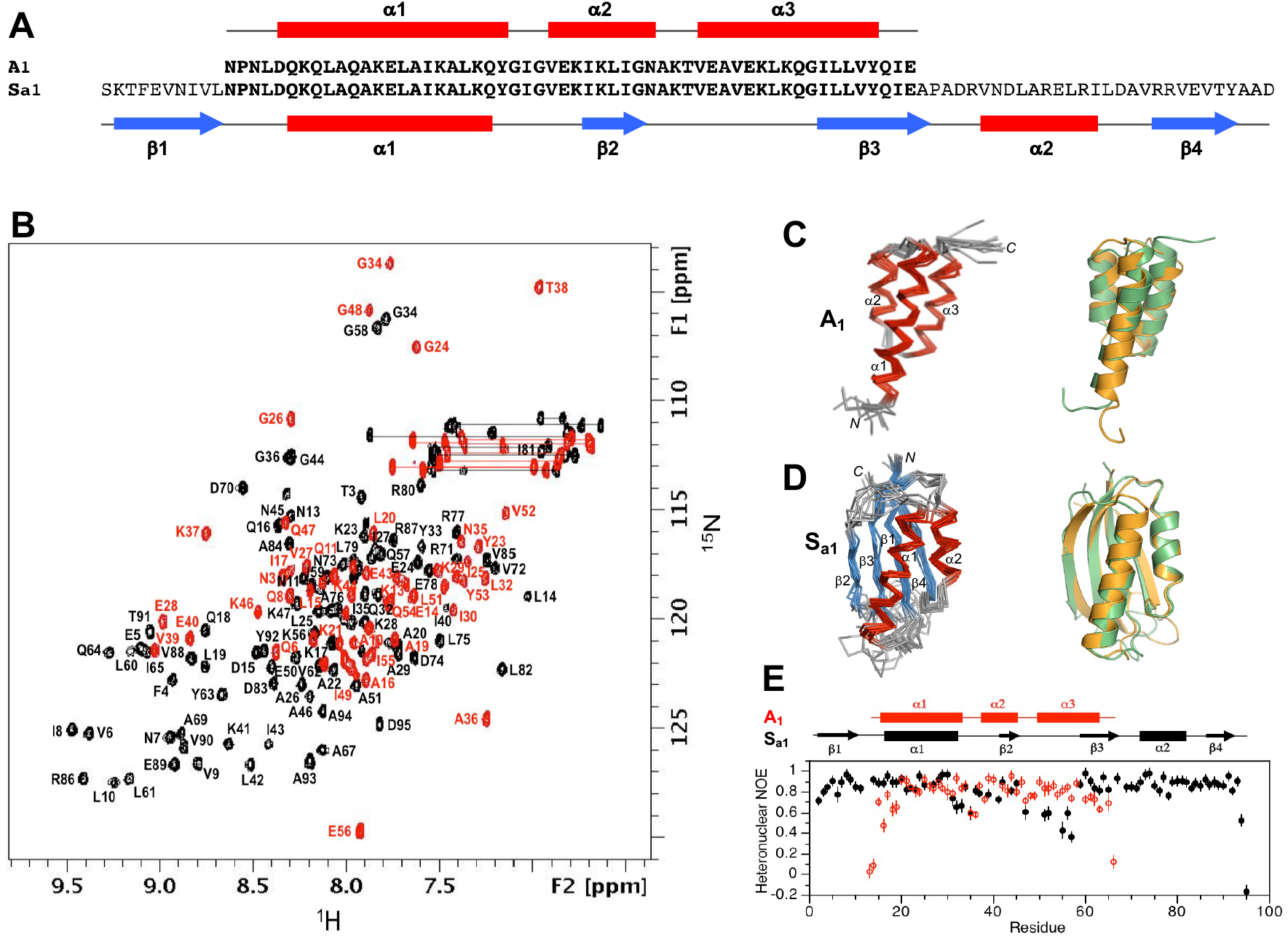
Structure and dynamics of A_1_ and S_a1_. (**A**) Sequence alignment of A_1_ and S_a1_, which are 100% identical over the 56 amino acid A-region. (**B**) Overlaid two dimensional ^1^H-^15^N HSQC spectra of S_a1_ (black) and A_1_ (red) with backbone amide assignments. Spectra were recorded at 25°C and 5°C, respectively. (**C**) Ensemble of 10 lowest energy CS-Rosetta structures for A_1_ (left panel). Superposition of the A_1_ structure (green) with the parent G_A_ fold (orange) (right panel). (**D**) Ensemble of 10 lowest energy CS-Rosetta structures for S_a1_ (left panel). Superposition of S_a1_ (green) with the parent S6 fold (orange) (right panel). (**E**) Backbone dynamics in designed proteins. Plot of {^1^H}-^15^N steady state heteronuclear NOE values at 600 MHz versus residue for A_1_ (red) and for S_a1_ (black). Error bars indicate ±1SD.

### Structural analysis of A_1_ and S_a1_

Overall, the 3α-helical bundle topology of A_1_ is very similar to the G_A_ parent structure from which it was derived ^46^. The sequence specific chemical shift assignments for A_1_ (**Fig. 3B**) were utilized to calculate a 3D structure with CS-Rosetta (**Fig. 3C, Table 1**). Our previous studies indicated close correspondence of CS-Rosetta and *de novo* structures for A- and B-folds with high sequence identity ^47^. The N-terminal residues 1-4 and the C-terminal residues 53-56 are disordered in the structure, consistent with {^1^H}-^15^N steady state heteronuclear NOE data (**Fig. 3E**). Likewise, S_a1_ has the same overall βαββαβ-topology as the parent S6 structure (**Fig. 3D, Table 2**). The backbone chemical shifts (**Fig. 3B**) were used in combination with main chain inter-proton NOEs (**Fig. S4**) to determine a three-dimensional structure utilizing CS-Rosetta (**PDB 7MN1)**. The conformational ensemble shows well-defined elements of secondary structure at residues 2-10 (β1), 16-32 (α1), 40-44 (β2), 59-67 (β3), 73-81 (α2) and 86-92 (β4). The principal difference from the native structure is that the β2-strand is seven amino acids shorter in S_a1_ than in S6. Heteronuclear NOE data show overall consistency with the structure, indicating that the long loop between the β2- and β3-strands from residues 45-58 is more flexible than other internal regions of the polypeptide chain (**Fig. 3E**).

**Table 1:** Structure statistics for A_1_, B_1_, and B_4_.

|  | A<sub>1</sub> | B<sub>1</sub> | B<sub>4</sub> |
| --- | --- | --- | --- |
| *A. Experimental chemical shift inputs* |  |  |  |
| <sup>13</sup>C<sub>α</sub> | 55 | 56 | 56 |
| <sup>13</sup>C<sub>β</sub> | 51 | 50 | 53 |
| <sup>13</sup>CO | 54 | 55 | 53 |
| <sup>15</sup>N | 54 | 55 | 54 |
| <sup>1</sup>H<sub>N</sub> | 54 | 55 | 54 |
| <sup>1</sup>H<sub>α</sub> | - | 55 | - |
| *B. RMSDs to the mean structure (Å)* |  |  |  |
| Over all residues |  |  |  |
| Backbone atoms | 1.73±0.47 | 0.86±0.22 | 0.85±0.34 |
| Heavy atoms | 2.27±0.53 | 1.37±0.29 | 1.43±0.45 |
| Secondary structures<sup>a</sup> |  |  |  |
| Backbone atoms | 0.90±0.29 | 0.67±0.22 | 0.62±0.25 |
| Heavy atoms | 1.55±0.42 | 1.12±0.25 | 1.33±0.41 |
| *C. Measures of structure quality (%)* |  |  |  |
| Ramachandran distribution |  |  |  |
| Most favored | 98.78±1.64 | 92.31±3.35 | 92.36±2.71 |
| Additionally allowed | 1.22±1.64 | 7.69±3.36 | 7.64±2.71 |
| Generously allowed | 0.00±0.00 | 0.00±0.00 | 0.00±0.00 |
| Disallowed | 0.00±0.00 | 0.00±0.00 | 0.00±0.00 |
| *D. Backbone RMSDs to the parent structure<sup>b</sup> (Å)* |  |  |  |
| Over all residues | 2.48 | 0.62 | 0.64 |
| Secondary structures | 1.21 | 0.57 | 0.49 |
| *E. PDB/BMRB codes* |  |  |  |
| PDBDEV | 00000083 | 00000084 | 00000085 |
| BMRB | 50907 | 50910 | 50909 |

<sup>a</sup> The secondary elements used were as follows: A<sub>1</sub>, residues 5-23, 27-35, 39-53; B<sub>1</sub>, residues 2-8, 13-19, 23-37, 43-46, 51-55; B<sub>4</sub>, residues: 2-8, 13-19, 23-37, 42-46, 51-55.
<sup>b</sup> The parent structure for A<sub>1</sub> is G<sub>A</sub> (PDB 2FS1). The parent structure for B<sub>1</sub> and B<sub>4</sub> is G<sub>B</sub> (PDB 1PGA). RMSDs were calculated by superimposing the mean structure from the NMR ensemble with either the mean structure (in the case of 2FS1) or the X-ray structure (in the case of 1PGA) for the parent.

**Table 2:**
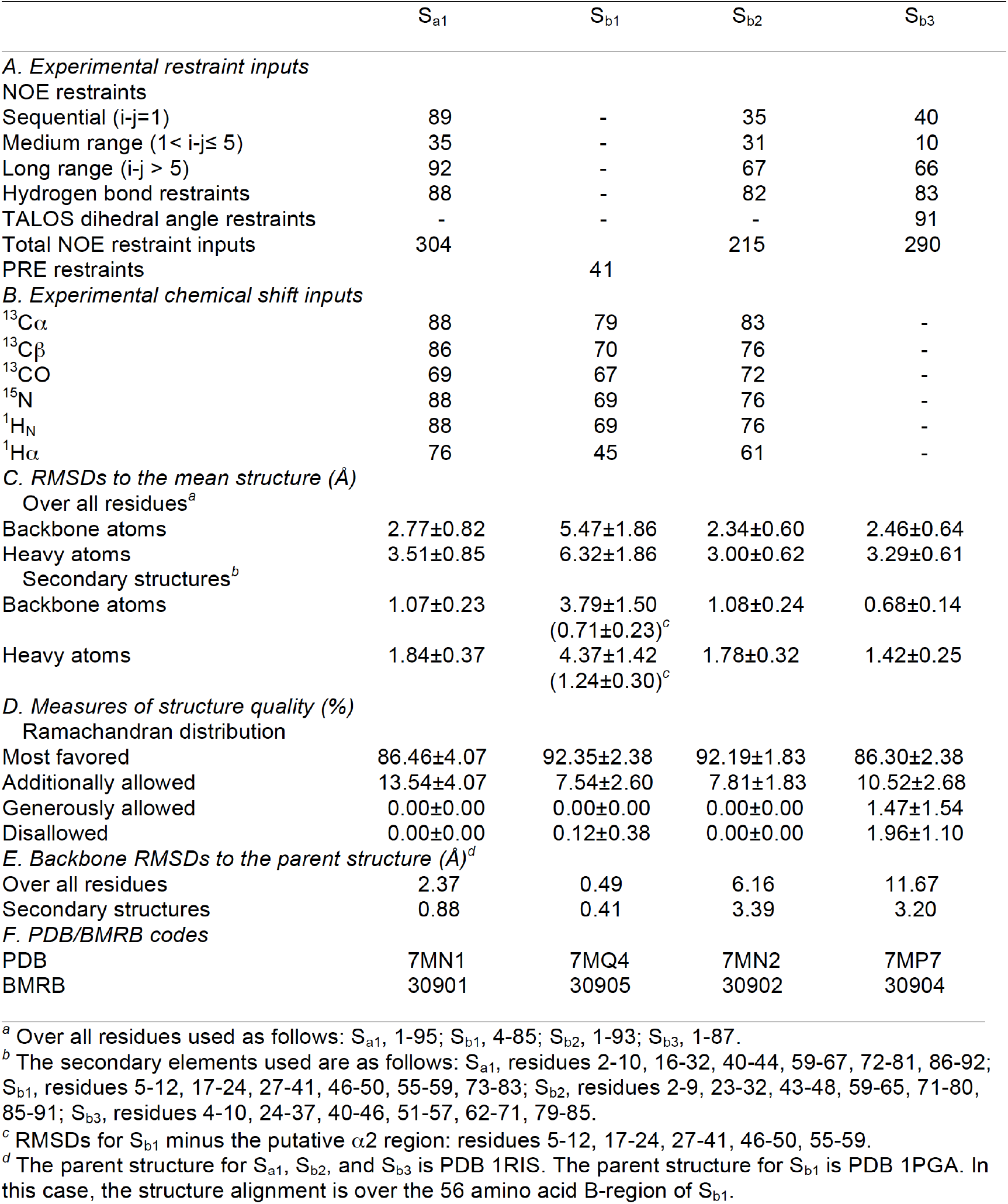
Structure statistics for S_a1_, S_b1_, S_b2_, and S_b3_.

|  | S<sub>a1</sub> | S<sub>b1</sub> | S<sub>b2</sub> | S<sub>b3</sub> |
| --- | --- | --- | --- | --- |
| *A. Experimental restraint inputs* |  |  |  |  |
| NOE restraints |  |  |  |  |
| Sequential (i-j=1) | 89 | - | 35 | 40 |
| Medium range (1 < i-j ≤ 5) | 35 | - | 31 | 10 |
| Long range (i-j > 5) | 92 | - | 67 | 66 |
| Hydrogen bond restraints | 88 | - | 82 | 83 |
| TALOS dihedral angle restraints | - | - | - | 91 |
| Total NOE restraint inputs | 304 |  | 215 | 290 |
| PRE restraints |  | 41 |  |  |
| *B. Experimental chemical shift inputs* |  |  |  |  |
| <sup>13</sup>C<sub>α</sub> | 88 | 79 | 83 | - |
| <sup>13</sup>C<sub>β</sub> | 86 | 70 | 76 | - |
| <sup>13</sup>CO | 69 | 67 | 72 | - |
| <sup>15</sup>N | 88 | 69 | 76 | - |
| <sup>1</sup>H<sub>N</sub> | 88 | 69 | 76 | - |
| <sup>1</sup>H<sub>α</sub> | 76 | 45 | 61 | - |
| *C. RMSDs to the mean structure (Å)* |  |  |  |  |
| Over all residues<sup>a</sup> |  |  |  |  |
| Backbone atoms | 2.77±0.82 | 5.47±1.86 | 2.34±0.60 | 2.46±0.64 |
| Heavy atoms | 3.51±0.85 | 6.32±1.86 | 3.00±0.62 | 3.29±0.61 |
| Secondary structures<sup>b</sup> |  |  |  |  |
| Backbone atoms | 1.07±0.23 | 3.79±1.50<br>(0.71±0.23)<sup>c</sup> | 1.08±0.24 | 0.68±0.14 |
| Heavy atoms | 1.84±0.37 | 4.37±1.42<br>(1.24±0.30)<sup>c</sup> | 1.78±0.32 | 1.42±0.25 |
| *D. Measures of structure quality (%)* |  |  |  |  |
| Ramachandran distribution |  |  |  |  |
| Most favored | 86.46±4.07 | 92.35±2.38 | 92.19±1.83 | 86.30±2.38 |
| Additionally allowed | 13.54±4.07 | 7.54±2.60 | 7.81±1.83 | 10.52±2.68 |
| Generously allowed | 0.00±0.00 | 0.00±0.00 | 0.00±0.00 | 1.47±1.54 |
| Disallowed | 0.00±0.00 | 0.12±0.38 | 0.00±0.00 | 1.96±1.10 |
| *E. Backbone RMSDs to the parent structure (Å)<sup>d</sup>* |  |  |  |  |
| Over all residues | 2.37 | 0.49 | 6.16 | 11.67 |
| Secondary structures | 0.88 | 0.41 | 3.39 | 3.20 |
| *F. PDB/BMRB codes* |  |  |  |  |
| PDB | 7MN1 | 7MQ4 | 7MN2 | 7MP7 |
| BMRB | 30901 | 30905 | 30902 | 30904 |

<sup>a</sup> Over all residues used as follows: S<sub>a1</sub>, 1-95; S<sub>b1</sub>, 4-85; S<sub>b2</sub>, 1-93; S<sub>b3</sub>, 1-87.
<sup>b</sup> The secondary elements used are as follows: S<sub>a1</sub>, residues 2-10, 16-32, 40-44, 59-67, 72-81, 86-92; S<sub>b1</sub>, residues 5-12, 17-24, 27-41, 46-50, 55-59, 73-83; S<sub>b2</sub>, residues 2-9, 23-32, 43-48, 59-65, 71-80, 85-91; S<sub>b3</sub>, residues 4-10, 24-37, 40-46, 51-57, 62-71, 79-85.
<sup>c</sup> RMSDs for S<sub>b1</sub> minus the putative α2 region: residues 5-12, 17-24, 27-41, 46-50, 55-59.
<sup>d</sup> The parent structure for S<sub>a1</sub>, S<sub>b2</sub>, and S<sub>b3</sub> is PDB 1RIS. The parent structure for S<sub>b1</sub> is PDB 1PGA. In this case, the structure alignment is over the 56 amino acid B-region of S<sub>b1</sub>.

### Comparison of A_1_ and S_a1_ structures

Although the 56 amino acid sequence of A_1_ is 100% identical to residues 11-66 of S_a1_, a significant fraction of the backbone undergoes changes between the two structures. Most notably, while the α1 helices in both A_1_ and S_a1_ are similar in length, the regions corresponding to the α2 and α3 helices of A_1_ form the β2 and β3 strands of S_a1_ (**Fig. 4A**). Core amino acids in the α1-helix of A_1_ correspond with residues that also contribute to the core of S_a1_. However, the α1-helix in S_a1_ contacts an almost entirely different set of residues (**Fig. 4B**). For example, amino acids L51, Y53, and I55 in the C-terminal tail of A_1_ do not have extensive contacts with α1 but the corresponding residues in S_a1_ (L61, Y63, and I65) form close core interactions with α1 as part of the β3-strand. Most of the other core residues contacting the α1-helix of S_a1_ are outside the 56 amino acid region coding for the A_1_ fold. These include F4, V6, I8, and L10 from the β1-strand; A67 from the β3-strand; V72, L75, and L79 from the α2-helix; and V85 from the loop between the α2-helix and the β4-strand. Two additional residues, V88 and V90 (β4) also contribute significantly to the core but do not contact α1. Thus, except for the original topological alignment of the α1-helices, the cores of the 3α and α/β-plait folds are largely non-overlapping. In total, approximately half of the residues participating in the S_a1_ core are not present in the A_1_ sequence.

**Fig. 4:**
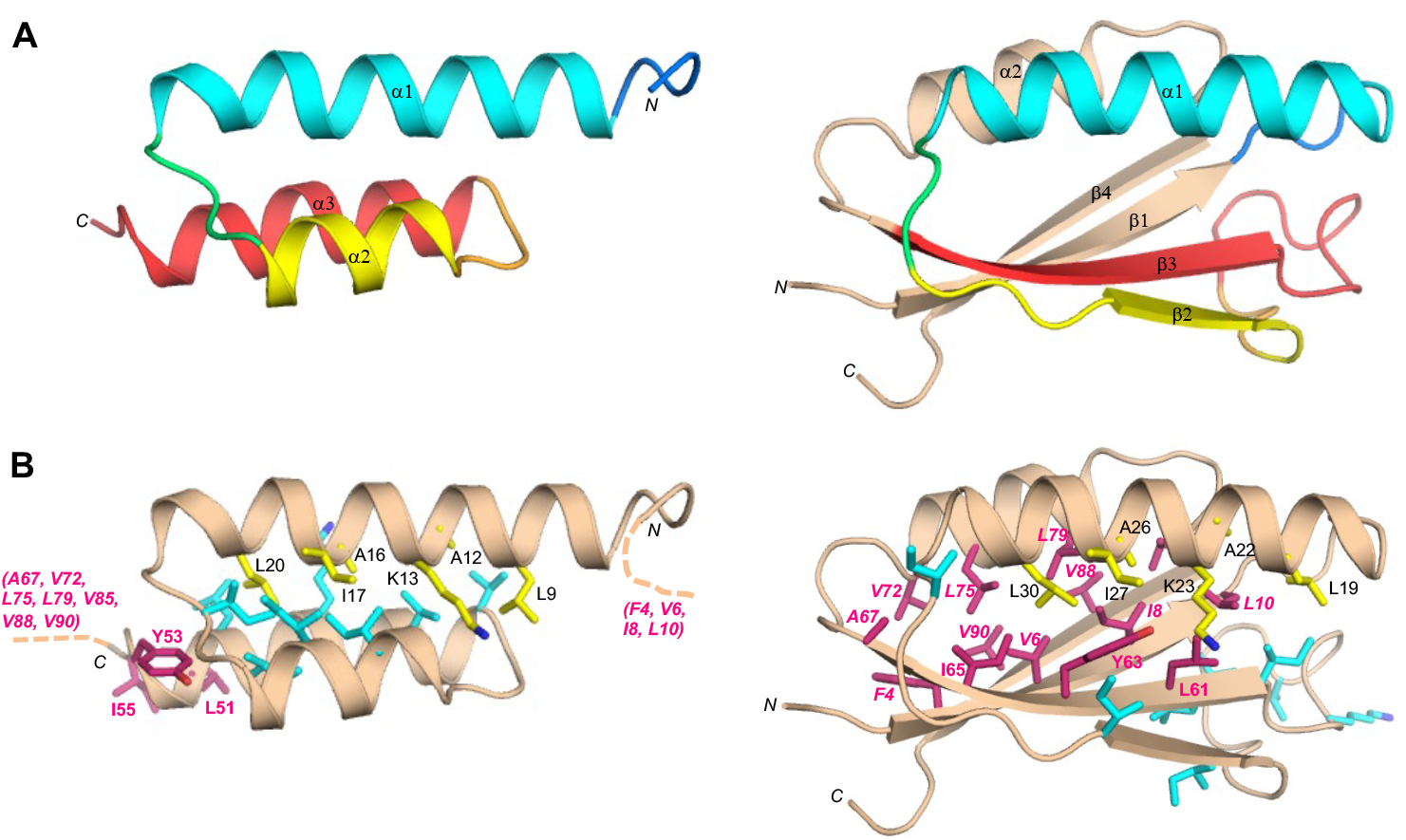
Structural differences between the 100% sequence identical regions of A_1_ and S_a1_. (**A**) Main chain comparisons. (Left panel) CS-Rosetta structure of A_1_ with color coding for secondary structured elements. (Right panel) Corresponding color-coded regions mapped onto the CS-Rosetta structure of S_a1_, illustrating changes in backbone conformation. Regions outside the 56 amino acid sequence of A_1_ are shown in wheat. (**B**) Side chain comparisons. (Left panel) Residues contributing to the core of A_1_ from the α1-helix (yellow), and from other regions (cyan). The non-a1 core residues from S_a1_ (pink) do not overlap with the A_1_ core (see text for further details). (Right panel) Residues contributing to the core of S_a1_ from the α1-helix (yellow), and most of the other participating core residues (pink). The non-a1 core residues from A_1_ are also shown (cyan), highlighting the low degree of overlap.

### Energetics of unfolding for A_1_/S_a1_

Far-UV CD spectra were measured for S_a1_ and A_1_ and their thermal unfolding profiles were determined by measuring ellipticity at 222nm versus temperature (**Fig. 5 and S5**). S_a1_ has a T_M_ of ^~^100°C and an estimated ΔG_folding_ of −5.3 kcal/mol at 25°C (**Fig. 5B**, **Table S1**) ^48^. The ΔG_folding_ of the parent S6 is −8.5 kcal/mol ^30^. The Rosetta energy of the S_a1_ design is similar to that of the native sequence (**Fig. S3**). A_1_ has a T_M_ of 65°C and a ΔG_folding_ = −4.0 kcal/mol at 25°C ^48^ (**Fig. 5A**, **Table S1**). The ΔG_folding_ of the parent G_A_ is −5.6 kcal/mol ^49,50^. The Rosetta energy of the A_1_ design is slightly more favorable than for the native sequence (**Fig. S3**).

**Fig. 5:**
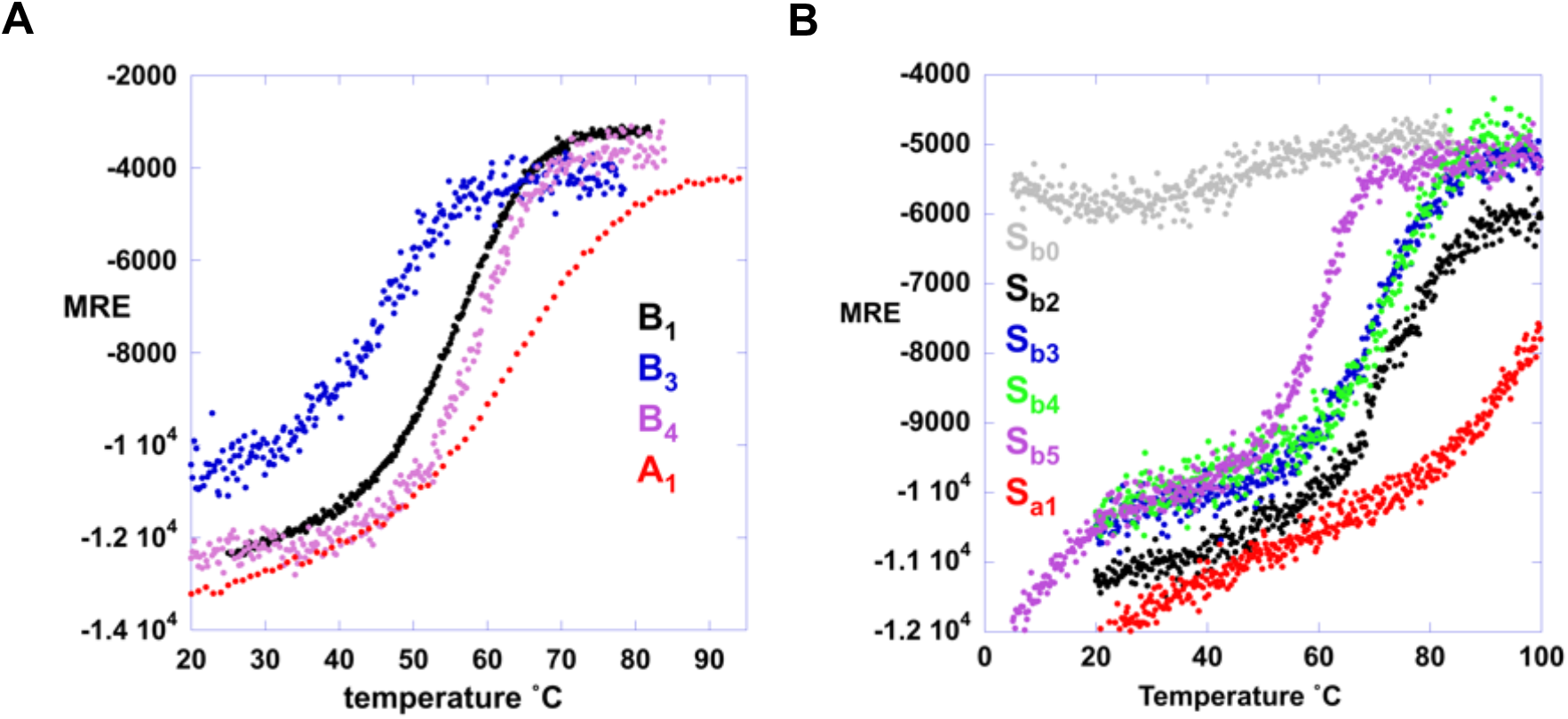
CD melting curves. (A) Ellipticity at 222nm plot versus temperature for A- and B- variations. (B) Ellipticity at 222nm plot versus temperature for S-variations. S_b0_ is a low-stability variant (F7V) of S_b1_ used to measure the temperature dependence of the unfolded state.

### HSA binding

Initial engineering of the fold switch was carried out without consideration of preserving function. As a result, A_1_ does not have detectible HSA binding affinity because two amino acids in the binding interface were mutated. Significant HSA-binding is recovered, however, when the surface mutations, E28Y and K29Y, are made in A_1_ (denoted A_2_). These mutations do not appear to affect the structure of A_1_ (**Fig. S5**) but result in HSA binding of K_D_ ≤ 1μM (**Table S1**). This was determined by measuring binding to immobilized HSA as described in Material and Methods.

### Protease inhibition

S_a1_ does not bind protease because C-terminal amino acids were not preserved in its design. It can be converted into a protease inhibitor, however, by replacing its three C-terminal amino acids (AAD) with DKLYRAL (denoted S_a1_I). A version of S_a1_I was also made that contains the exact 56 amino acid A_2_ sequence by making E38Y, K39Y mutations (denoted Sa2I). S_a1_, S_a1_I, and Sa2I are similar in structure by CD analysis (**Fig. S5**). The inhibition constant of Sa2I with the engineered subtilisin was determined to be 50nM as described in Materials and Methods (**Table S1**). Thus, a stable A-fold with HSA-binding function can be embedded within a 99 amino acid S-fold with protease inhibitor function (**Fig. 2C and E**). It should be noted that all HSA contact amino acids are preserved in both the A_2_ and Sa2I sequences, but the three-dimensional topology necessary to form the HSA contact surface occurs only in the A-fold. Nevertheless, Sa2I was observed to bind weakly to HSA (K_D_ ^~^100μM), **Table S1)**. This weak affinity suggests that some Sa2I molecules may populate the 3α fold even though the α/β-plait fold strongly predominates.

### Designing a switch from α/β-plait protease inhibitor to β–grasp IgG-binding protein

In designing an S- to B-fold switch, we used two topological alignments. The first was between SI- and B-folds, where the β1 strands of each fold were aligned (**Fig. S2B, Fig. S6A**). The second alignment was between S’I- and B-folds, where the long loop between β2 and β3 in SI was shortened in S’I to be more consistent with natural protease inhibitors. In this scheme, the α1β3β4 topology of the B-fold was aligned with the α1β2β3 topology of the S’I-fold (**Fig. 6A, Fig. S2C**).

**Fig. 6:**
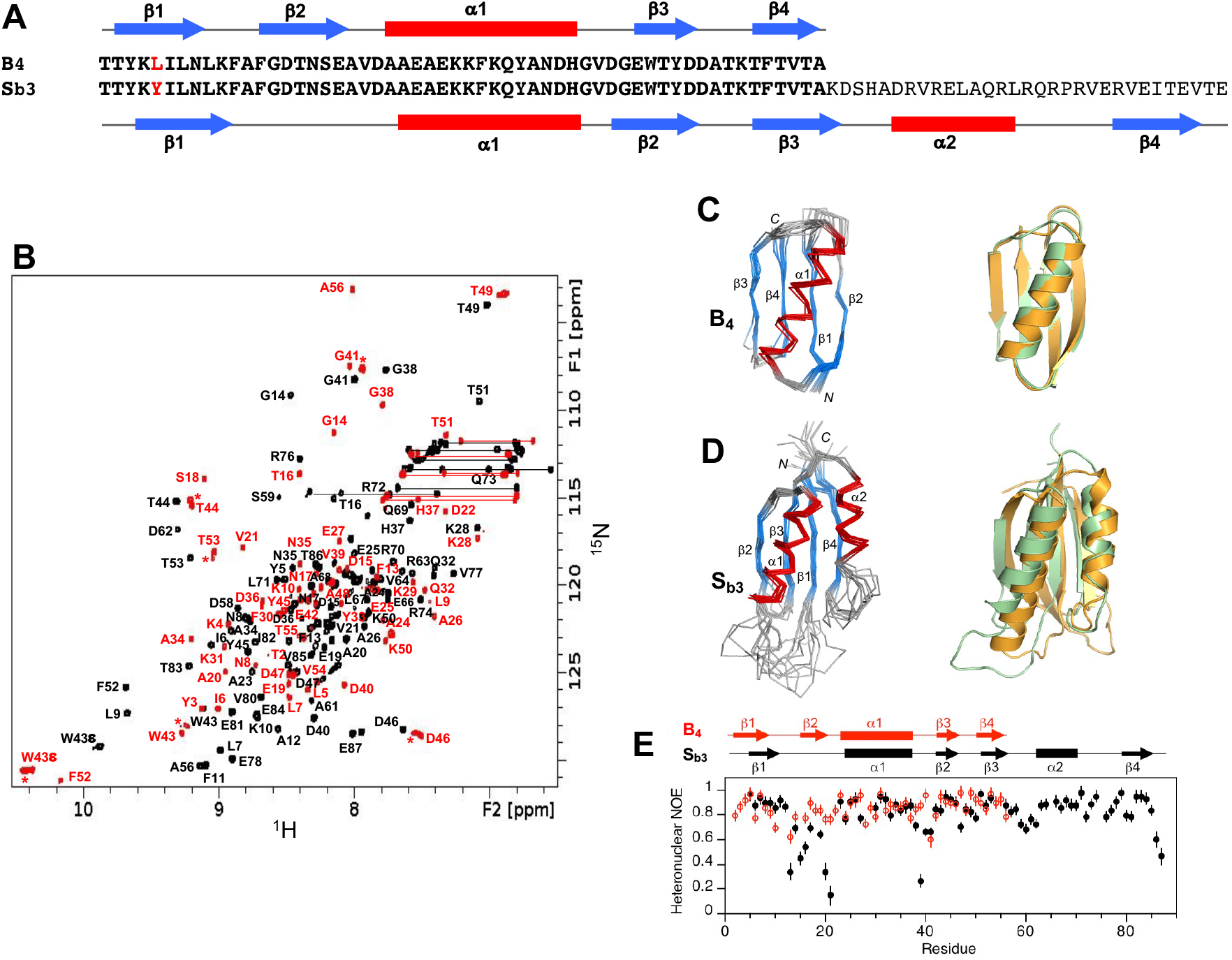
Structure and dynamics of S_b3_ and B_4_. (**A**) Sequence alignment of B_4_ and S_b3_, differing by one residue (L5Y) over the 56 amino acid B-region. (**B**) Overlaid two dimensional ^1^H-^15^N HSQC spectra of S_b3_ (black) and B_4_ (red) with backbone amide assignments. Spectra were recorded at 25°C. The A56 peak is an aliased signal. Peaks labeled with an asterisk decrease in relative intensity as the B_4_ concentration is lowered, indicating the presence of a weakly associated putative dimer in addition to monomer. (**C**) Ensemble of 10 lowest energy CS-Rosetta structures for B_4_ (left panel). Superposition of the B_4_ structure (green) with the parent G_B_ fold (orange) (right panel). (**D**) Ensemble of 10 lowest energy CS-Rosetta structures for S_b3_ (left panel). Superposition of S_b3_ (green) with the parent S6 fold (orange) (right panel). (**E**) Plot of {^1^H}-^15^N steady state heteronuclear NOE values at 600 MHz versus residue for B_4_ (red) and S_b3_ (black). Error bars indicate ±1SD.

### Design and characterization of B_1_, S_b1_, B_2_, and S_b2_

In the first approach, alignment of the β1-strands of the B-fold and the S-fold and subsequent mutation to resolve catastrophic interactions produced low energy switch candidates denoted B_1_ and S_b1_. The exact sequence of B_1_ is embedded in S_b1_ at positions 4-59 (**Fig. S6A**). The computational models of B_1_ and S_b1_ show relatively small increases in energy compared to the corresponding relaxed native structures (**Fig. S3**). The NMR structure of B_1_ displayed a ββαββ topology identical to that of the parent B-fold, with a backbone RMSD of ^~^0.6 Å (**Fig. S6B, C**). The topology of S_b1_ is not the same as the parent S6 structure, however, and instead has a fold similar to that of B_1_ (**Fig. S6B, D**; **Fig. S7, PDB 7MQ4**). Introducing 13 mutations into S_b1_ generated a protein denoted S_b2_ (**Fig. S8**). S_b2_ contains four β-strands and two α-helices and has the general features of the parent S-fold (**Fig. S9, PDB 7MN2**). The 56 amino acid version of S_b2_ (denoted B_2_) has a significantly higher Rosetta energy than B_1_, however, and is presumably unfolded (**Fig. S3**). Thus, neither the B_1_/S_b1_ nor B_2_/S_b2_ protein pairs resulted in high identity sequences with different folds. Nonetheless, B_1_ is 80% identical to the corresponding embedded region in the S-folded protein S_b2_ (**Fig. S9A**). The structures of B_1_, S_b1_, and S_b2_ are described further in the Supplement and Tables 1 and 2.

### Design of S_b3_ and B_3_

To improve the design of the S to B switch we aligned the B-fold with the S’ inhibitor fold and chose an alignment that creates a topological match between α1β3β4 in B and α1β2β3 in S’ (**Fig. S2C**). Mutation to resolve deleterious interactions in this alignment produced low energy switch candidates denoted B_3_ and S_b3_ **(Fig. S10)**. The exact sequence of B_3_ is embedded in S_b3_ at positions 1-56. The energy of the computational model for S_b3_ is slightly more favorable than the relaxed native structure. The designed model of B_3_ shows relatively small increases in energy compared to the relaxed native structure (**Fig. S3**).

### Structural analysis of S_b3_ and B_3_

NMR-based structure determination indicated that S_b3_ has a βαββαβ secondary structure and an S-fold topology (**Fig. 6A,B,D, PDB 7MP7**). Ordered regions correspond with residues 4-10 (β1), 24-37 (α1), 42-46 (β2), 51-56 (β3), 62-70 (α2), and 79-85 (β4). Comparison of S_b3_ with the parent S-fold indicates that the β1/α2/β4 portion of the fold is similar in both. In contrast, the β1-α1 loop is longer in S_b3_ (13 residues) than in the parent S-fold (5 residues), while α1, β2, the β2-β3 loop, and β3 are all shorter than in the parent (**Fig. 6D**). Consistent with the S_b3_ structure, the 13 amino acid β1-α1 loop is highly flexible (**Fig. 6E**). We also expressed and purified a truncated protein corresponding to the embedded B-fold, the 56 amino acid version of S_b3_ (denoted B_3_). The 2D ^1^H-^15^N HSQC spectrum of B_3_ at 5°C and low concentrations (<20 μM) was consistent with a predominant, monomeric B-fold (**Fig. S11**) but showed significant exchange broadening at 25°C, indicative of low stability (see below). Presumably the low stability is due to less favorable packing of Y5 in the core of the B-fold compared with a smaller aliphatic leucine. However, additional, putatively oligomeric, species were also present for which relative peak intensities increased with increasing protein concentration. Due to its relatively low stability and sample heterogeneity, B_3_ was not analyzed further structurally.

### Design and analysis of point mutations that switch the fold of S_b3_

We used the NMR structure of S_b3_ to design a point mutation, tyrosine 5 to leucine (Y5L), that would stabilize the embedded B-fold without compromising native contacts in the S-fold (**Fig. S10**). This mutant was therefore expected to shift the population to the B-fold. The Y5L mutant of S_b3_ (denoted S_b4_) exhibited approximately twice the number of amide cross-peaks in the HSQC spectrum relative to the S_b3_ sample. Comparison of the spectrum of S_b4_ with spectra of S- and B-folds for closely related sequences indicated that S_b4_ populates both S- and B-states simultaneously in an approximately 1:1 ratio at 25°C (**Fig. S12**). A complete NMR analysis of the kinetics of interconversion will be reported elsewhere. The Y5L mutation also was introduced into B_3_ to determine its effect on the stability of the B-fold in the 56 amino acid protein. The new protein, denoted B_4_, is indeed more stable than B_3_ (**Fig. 5A, Table S1**). Assignment and structure determination of B_4_ showed its topology to be identical to the parent B-fold (**Fig. 6B, C**). At concentrations above 100 μM, B_4_ displayed a tendency for weak self-association similar to that seen for B_3_.

Finally, we designed a mutation of leucine 67 to arginine (L67R) in S_b4_ to destabilize the S-fold without changing the sequence of the embedded B-fold. The mutant is denoted as S_b5_ (**Fig. S10**). This was expected to shift the population to the B-fold. The 2D ^1^H-^15^N HSQC spectrum of S_b5_ indicates that the L67R mutation does indeed destabilize the S-fold, with the loss of S-type amide cross-peaks and the concurrent appearance of a new set of signals indicating a switch to a B-fold. Superposition of the spectrum of S_b5_ with that of B_4_ shows that the new signals in S_b5_ largely correspond with the spectrum of B_4_ (**Fig. S13**). Thus, the L67R mutation shifts the equilibrium from the S-fold to the B-fold. The additional signals (^~^25-30) in the central region of the HSQC spectrum that are not detected in B_4_ are presumably due to the disordered C-terminal tail of S_b5_. The C-terminal tail of S_b5_ does not appear to interact extensively with the B-fold, as evidenced by few changes in chemical shifts or peak intensities in the B-region of S_b5_ compared with B_4_.

### Structural comparison of S_b3_ and B_4_

The aligned amino acids 1-56 of S_b3_ and B_4_ have 98% sequence identity, the only difference being an L5Y mutation in S_b3_ (**Fig. 6A**). The global folds of S_b3_ and B_4_ have large-scale differences, however (**Fig. 7A, Fig. S4**). The β1-strands, while similar in length, are in opposite directions in S_b3_ and B_4_. The β1-strand forms a parallel stranded interaction with β4 in B_4_, but an antiparallel interaction with the corresponding β3-strand in S_b3_. Whereas residues 9-20 form the 6-residue β1-β2 turn and the 6-residue β2-strand of B_4_, these same amino acids constitute the end of β1 and 10 residues of the large disordered β1-α1 loop in S_b3_. The remainder of the B-region is topologically similar, with the α1/β3/β4 structure in B_4_ matching the α1/β2/β3 structure in S_b3_. Overall, however, the order of H-bonding in the 4-stranded β-sheets is quite different, with β2β3β1β4 in S_b3_ and β3β4β1β2 in B_4_.

**Figure 7:**
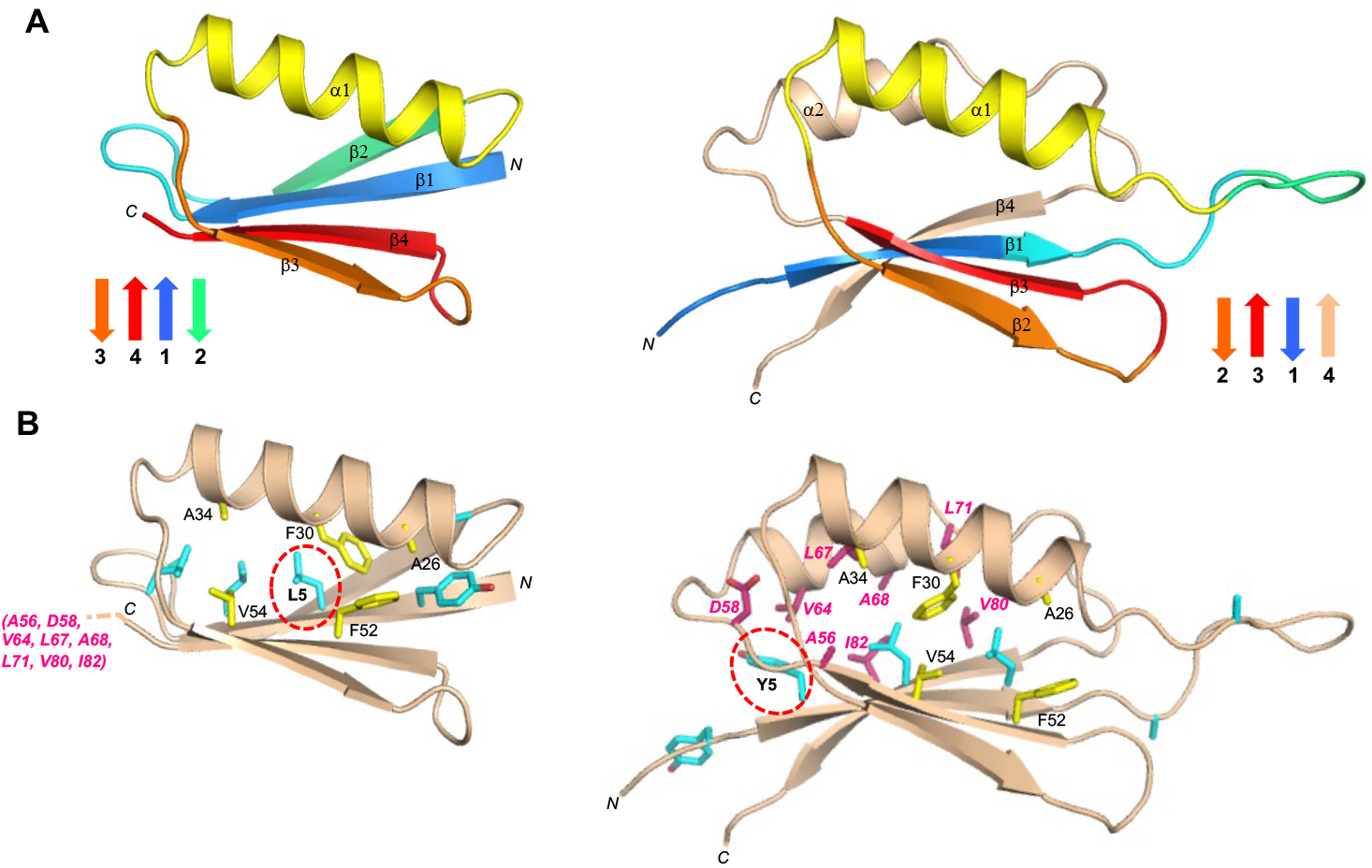
Structural differences in the high (^~^98%) sequence identity regions of B_4_ and S_b3_. (**A**) Main chain comparisons. (Left panel) CS-Rosetta structure of B_4_ with secondary structure elements color coded. (Right panel) Corresponding color-coded regions mapped onto the CS-Rosetta structure of S_b3_, showing changes in backbone conformation. Regions outside the 56 amino acid sequence of B_4_ are shown in wheat. (**B**) Side chain comparisons. (Left panel) Residues contributing to the core of B_4_ from α1/β3/β4 (yellow), and from other regions (cyan). The non-α1/β2/β3 core residues from S_b3_ (pink) do not overlap with the B_4_ core (see text for further details). (Right panel) Residues contributing to the core of S_b3_ from α1/β2/β3 (yellow), and the other participating core residues (pink). The non-α1/β2/β3 core residues from B_4_ are also shown (cyan).. The single L5Y amino acid difference between B_4_ and S_b3_ is highlighted.

The main core residues of B_4_ consist of Y3, L5, L7, and L9 from β1, A26, F30, and A34 from α1, and F52 and V54 from β4 (**Fig. 7B**). In S_b3_, the topologically equivalent regions of the core are A26, F30, and A34 from α1, and F52 and V54 from β3. Residues Y5, L7, and L9 from the β1 strand of S_b3_ also form part of the core, but with different packing from B_4_ due to the reverse orientation of β1. Residues A12 and A20, which contribute to the periphery of the core in B_4_, are solvent accessible in the β1-α1 loop of S_b3_. Most of the remaining core residues of S_b3_ come from outside of the B-region and include amino acids from β3 (A56), α2 (V64, L67, A68, L71), and β4 (V80 and I82).

### Energetics of unfolding for B_3_/S_b3_, B_4_/S_b4_, and S_b5_

Far-UV CD spectra were measured for B_3_, B_4_, S_b3_, S_b4_, and S_b5_ and their thermal unfolding profiles were determined by measuring ellipticity at 222nm versus temperature (**Fig. 5, S10, Table S1**). As described above, the predominant form of S_b3_ is an S-fold. CD and NMR analysis show that B_3_ is predominantly a B-fold with a ΔG_folding_ of −1.2 kcal/mol at 25°C ^48^. From the NMR analysis, it appears that the B-fold is in equilibrium with putatively dimeric states. This creates a situation in which the B-fold is both temperature dependent and concentration dependent. The predominant form at 5°C and ≤18μM is the B-fold, however. The low stability and concentration-dependent behavior of B_3_ may indicate that some propensity for S-type conformations could persist in the 56-residue protein.

S_b4_ has a temperature unfolding profile very similar to S_b3_ even though both S- and Bare equally populated at 25°C in S_b4_ (**Fig. 5**). This shows that the Y5L mutation results in two folds that are almost isoenergetic and both thermodynamically stable relative to the unfolded state. Further, because S- and B-folds are in equilibrium and equally populated, the free energy of switching to the B-fold from the S-fold (ΔG_B-fold/S-fold_) is ^~^ 0 kcal/mol at 25°C. The switch equilibrium reflects the influence of the antagonistic B-fold on the S-fold population in S_b4_, where the leucine at residue 5 helps stabilize the alternative B-state at the expense of the S-state. Thermal denaturation by CD shows that B_4_ has a ΔG_folding_ = −4.1 kcal/mol at 25°C ^48^. The thermal unfolding profile of S_b5_ shows a low temperature transition with a midpoint ^~^10°C and a major transition with a midpoint of ^~^60°C (**Fig. 5B**). The NMR analysis indicates that the major transition is unfolding of the B-fold. Thus, the arginine at 67 in S_b5_ makes the B-fold more favorable by making the S-fold unfavorable, consistent with the change in population from mixed to B-fold observed by NMR.

### Protease inhibition

The S_b3_ protein is closely related to S’I but lacks inhibitor function because C-terminal amino acids were changed in the design of the switch. It can be converted into a protease inhibitor, however, by altering C-terminal amino acids VTE to DKLYRAL. This mutant is denoted S_b3_I. S_b3_ and S_b3_I appear similar in structure by CD analysis (**Fig. S10**). The K_I_ for S_b3_I with subtilisin was determined to be 50nM (**Table S1**).

### IgG binding

Binding to IgG was determined for B_3_ and S_b3_I (**Table S1**). B_3_ and S_b3_I bound to IgG Sepharose with K_D_ ≤ 1μM and 10μM, respectively. Presumably S_b3_I has significant IgG-binding activity because the α1β3 IgG binding surface of the B-fold is largely preserved in the S-fold. Thus, S_b3_I is a dual-function protein with both IgG-binding and protease inhibitor functions (**Fig. 2F**).

## DISCUSSION

The entire network of intersecting pathways between the S-, A-, and B-folds is summarized in **Fig. 8**. The first node on the pathway is a functional switch from RNA binding protein to protease inhibitor without a fold switch. The α/β plait is a common fold, and proteins with this basic topology include many different functions ^32^. Engineering the SI and S’I nodes illustrates how protease inhibitor function can arise in the α/β plait topology with a few mutations. Replacing only C-terminal amino acids in the S6 protein creates interaction with the substrate binding cleft of the protease (**Fig. 2A, B**). This C-terminal interaction plus adventitious contact between the β–sheet surface of the α/β plait and two α–helices in the protease results in protease inhibition in the 50nM range. Based on the structure of S6 in the 30S complex, the C-terminal modification may not have major effects on binding interactions with ribosomal RNA and the S15 protein (**Fig. 2A**). Thus, the transition from RNA binding protein to protease inhibitor likely is uninterrupted. An insertion in the β1-α1 loop and a deletion β2-β3 loop in the SI-inhibitor creates a topology that more closely resembles natural prodomain-type inhibitors^36,38,51^ and creates an α1β2β3 motif in the S’-fold that is similar to the α1β3β4 motif of the B-fold. This topological similarity brings the S’I closer to an intersection with the B-fold. Thus, SI and S’I nodes are both a functional switch and a branch point for switching the S-fold into the A- and B-folds, respectively.

**Figure 8:**
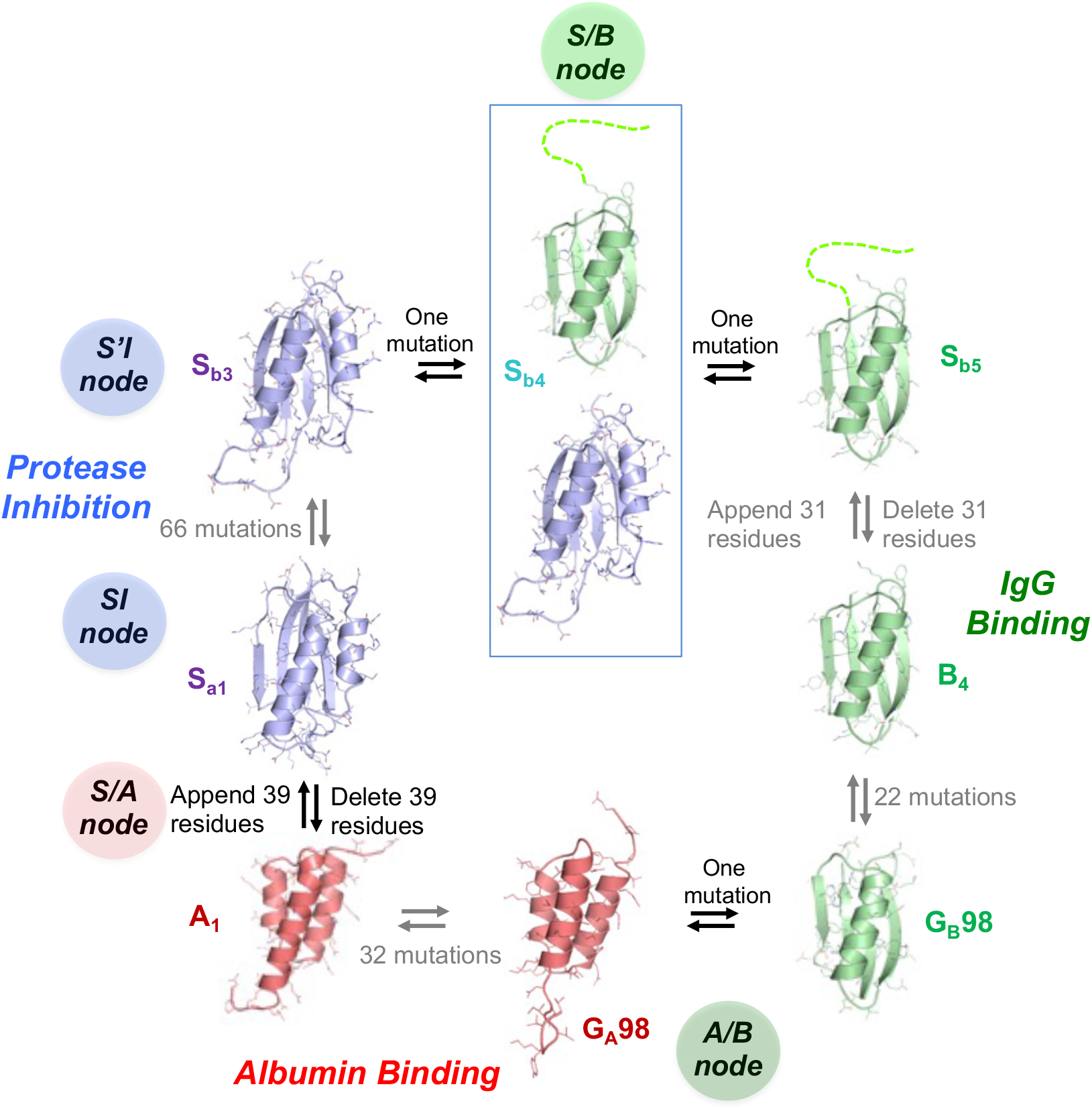
Sequence-fold relationships of engineered S/A, S/B, and A/B nodes. Switches between stable folds can be induced by single amino acid mutation or deleting/appending terminal sequence that stabilizes the S-fold. Blue denotes an S-fold, green a B-fold, and red an A-fold. Gray arrows connect proteins that have been reengineered without a fold switch. S_b4_ is observed with two folds simultaneously. The G_A_98 and G_B_98 structures are from PDB codes 2LHC and 2LHD, respectively.

Engineering nodes at fold intersections required designing sequences that are compatible with native interactions in two different folds. We used simple rules to do this. The first rule was to align topologies rather than maximizing sequence similarities. Identifying a common topology can help determine a register that has fewer irreconcilable clashes. For example, topological alignment of the α1 helix of the SI fold and the α1 helix of the A-fold facilitated engineering the fold switch because the regions flanking α1 of the SI-fold proved to be capable of encoding two different fold motifs. When topological alignment is poor, as was the case with S- and B-folds, it was helpful to look for natural variations in the turns of the longer fold to create better alignment. Variation in loops and turns in a larger fold creates more freedom of design and higher probability of switches. Once an alignment is chosen, the basic rule in resolving catastrophic clashes is to conserve original amino acids when possible. This reduces the uncertainties involved in computational design. The Rosetta energy function was not used to predict a favorable alignment but was important in evaluating mutations to resolve clashes once an alignment was chosen.

Selecting mutations compatible with two sets of native interactions required tradeoffs in the native state energetics of each individual fold to gain sequence identity ^2,8^. A node may be produced in cases in which both alternative folds are stable relative to the unfolded state. Stability relative to the unfolded state (i.e. a state with little secondary structure) was determined by CD melting (**Fig. 5**). It was informative to examine the stability of both short (56 residue) and longer forms of a putative node sequence. The independent stability of the G-fold can be determined in the short form without the antagonism from the S-fold that is present in the longer sequence. The stabilities of the A_1_ and A_2_ proteins are about −4 kcal/mol at 25°C ^48^ compared to −5.6 kcal/mol for the native G_A_ protein ^46^. The stabilities of B_3_ and B_4_ are −1.2 and - 4.1 kcal/mol, respectively, at 25°C ^48^ compared to −6.7 kcal/mol for the native G_B_ protein ^52^. For the longer sequences, the ΔG_folding_ of S_a1_ and S_b3_ are −5.3 kcal/mol and −3.5 kcal/mol, respectively, at 25°C ^48^ compared to −8.5 kcal/mol for the native S6 protein ^30^.

In the case of the S-folds, however, the energetic effects of the stable, embedded G-fold must also be considered. Since the equilibria between both folded states and the unfolded state are thermodynamically linked, the free energy of a switch to a G-fold from an S-fold (ΔG_G-fold/S-fold_) is approximated by the difference in ΔG_folding_ (ΔΔG_folding_) between the short and long forms of a node protein. For example, based on ΔG_folding_ for A_1_ and S_a1_, the predicted ΔG_A-fold/S-fold_ of S_a1_ is 1.3kcal/mol. This is consistent with the structure of the predominant S-fold determined by NMR but also with the small population of 3α fold suggested by weak HSA-binding. From the thermal denaturation profiles of B_3_ and S_b3_, the predicted ΔG_B-fold/S-fold_ of S_b3_ is 2.3kcal/mol, a value consistent with the stable S-fold observed in NMR experiments. The S_b3_ sequence is also approaching a critical point, however. A substitution in S_b3_ that stabilizes the B-fold (Y5L) shifts the equilibrium of S_b4_ to an approximately equal mixture of B- and S-folds. That is, ΔG_B-fold/S-fold_ of S_b4_ is ^~^ 0 kcal/mol at 25°C. One further substitution that destabilizes the S-fold (L67R) shifts the population of S_b5_ to a stable B-fold (ΔG_B-fold/S-fold_ ≤ −5 kcal/mol) (**Fig. 8**).

The existence of nodes between folds has implications for the evolution of new functions. In the case of the S/A node, all contact amino acids for HSA exist within the S-fold of the protease inhibitor Sa2I albeit in a cryptic topology. Deletion of amino acids 67-99 (A_2_) results in loss of inhibitor function and a fold switch from α/β plait to 3α. Acquisition of HSA binding activity (K_D_ < 1μM) results from unmasking the cryptic HSA binding amino acids via the fold switch (**Fig. 2E**). This level of binding affinity could be biologically relevant since the concentration of HSA in serum is > 500μM^53^. In the case of the S/B node, the α1β3 motif contains all IgG contact amino acids and S_b3_I has some affinity for both IgG (K_D_ = 10μM) and protease (K_I_ = 50nM). In this case, the Y5L mutation (S_b4_) or a deletion of 57-91 (B_4_) causes a fold switch from α/β plait to the β–grasp and results in tighter IgG binding (K_D_ ≤1μM) (**Fig. 2F**). This level of binding affinity could also be biologically relevant since the concentration of IgG in serum is >50μM (or >100μM Fc binding sites) ^54^. We have previously shown that an A-fold with HSA binding function can be switched to a B-fold with IgG-binding function via single amino acid substitutions that switch the folds and unmask cryptic contact amino acids for the two ligands ^34,35^.

In conclusion, it was possible to connect three common folds in a network of high-identity nodes that form critical points between two folds. As in other complex systems, a small change in a protein near a critical point can have a “butterfly effect” on how the folds are populated. This property of the protein folding code means that proteins with multiple folds and functions can exist in highly identical amino acid sequences. This suggests that evolution of new folds and functions sometimes can follow uninterrupted mutational pathways.

## MATERIALS AND METHODS

### Mutagenesis, protein expression and purification

Mutagenesis was carried out using Q5^®^ Site-Directed Mutagenesis Kits (NEB). G_A_ and G_B_ variants were cloned into a vector (pA-YRGL) encoding the sequence:

~~~
MEAVDANSLAQAKEAAIKELKQYGIGDKYIKLINNAKTVEGVESLKNEILKALPTEGSGEEDKQYRGL-
~~~

as an N-terminal fusion domain ^46^. Cell growth was carried out by auto-induction ^34,55^. Cells were harvested by centrifugation at 3,750 x g for 20 minutes and lysed by sonication on ice in 0.1M KPO_4_, pH 7.2. Cellular debris were pelleted by centrifugation at 10,000 x g for 15 minutes. Supernatant was clarified by centrifugation at 45,000 x g for 30 minutes. Proteins were purified using a second generation of the affinity-cleavage tag system employed previously to purify switch proteins ^34,56^. The second-generation tag (*YRGL-tag*) results in high-level soluble expression of the switch proteins and also enables capture of the fusion protein by binding tightly to an immobilized processing protease via the C-terminal EEDQYRGL sequence. Loading and washing were at 5 mL/min for a 5mL *Im-Prot* column using a running buffer of 20mM KPO_4_, pH 6.8. The amount of washing required for high purity depends on stickiness of the target protein and how much of it is bound to the column. We typically wash with 10 column volumes (CV) of wash solution followed by 3 CV 0.5M NaCl and then ^~^ 10 CV running buffer. This can be repeated as necessary. The 0.5M NaCl shots are repeated until the amount of absorbance release with each high-salt shot becomes small and constant. All the high-salt solution is washed out before initiating the cleavage. The target protein was cleaved from the *Im-Prot* column by injecting 15 mL of imidazole solution (0.1 mM) at 1mL/min, 22°C. The cleaved protein typically elutes as a sharp peak in 2-3 CV. The purified protein was then concentrated to 0.2 to 0.3 mM, as required for NMR analysis. The columns were regenerated by injecting 15 mL of 0.1 N H_3_PO_4_ (0.227 mL concentrated phosphoric acid (85%) per 100ml) at a flow rate of ^~^ 1 CV/min. The wash solution was neutralized immediately after stripping. The purification system is available from Potomac Affinity Proteins.

Protease inhibitor proteins were purified by binding to *Im-Prot* media and then stripping off the purified inhibitor with 0.1N H_3_PO_4_. Samples were then immediately neutralized by adding 1/10 volume 1M K_2_HPO_4_.

### Rosetta calculations

Rosetta energies of all designed structures were generated using the Slow Relax routine ^44^. 1000 decoys were calculated for each design. PDB coordinates and energy parameters for the lowest energy decoy for each design are included as supplemental files.

### Circular Dichroism (CD)

CD measurements were performed in 100mM KPO_4_, pH 7.2 with a Jasco spectropolarimeter, model J-1100 with a Peltier temperature controller. Quartz cells with path lengths of 0.1 cm and 1cm were used for protein concentrations of 3 and 30 μM, respectively. The ellipticity results were expressed as mean residue ellipticity, [θ], deg cm^2^ dmol^−1^. Ellipticities at 222 nm were continuously monitored at a scanning rate of 0.5 deg/min. Reversibility of denaturation was confirmed by comparing the CD spectra at 20°C before melting and after heating to 100°C and cooling to 20°C.

### Measuring HSA and IgG binding affinity

Affinity of proteins to HSA and IgG was determined by their retention on immobilized ligand. HSA and rabbit IgG were immobilized by reaction with NHS-activated Sepharose 4 Fast Flow (Cytiva) according to the manufacturer’s instructions. The concentration of immobilized HSA was 100μM. The concentration of immobilized IgG was 50μM (i.e. 100μM Fc binding sites). Generally 0.2mL of a 5μM solution of the test protein was injected onto a 5mL column at a flow rate of 0.5mL/min. Determination of binding affinity assumes that binding is in rapid equilibrium such that the elution volume is proportional to the fraction of test protein bound to 100μM of binding sites. Proteins that are completely retained after 20 column volumes (CV) are assessed to have K_D_ ≤ 1μM. Completely retained proteins are stripped from the column with 0.1N H_3_PO_4_ at the end of the run.

### Measuring protease inhibition

Competitive inhibition constants (K_I_) are measured by determining the K_M(apparent)_ for the substrate QEEYSAM-AMC and RASProtease(I)^41^ in the presence of 100nM test protein.

### NMR Spectroscopy

Isotope-labeled samples were prepared at 0.2-0.3 mM concentrations in 100 mM potassium phosphate buffer (pH 7.0) containing 5% D_2_O. NMR spectra were collected on Bruker AVANCE III 600 and 900 MHz spectrometers fitted with Z-gradient ^1^H/^13^C/^15^N triple resonance cryoprobes. Standard double and triple resonance experiments (HNCACB, CBCA(CO)NH, HNCO, HN(CA)CO, and HNHA) were utilized to determine main chain NMR assignments. Inter-proton distances were obtained from 3D ^15^N-edited NOESY and 3D ^13^C-edited NOESY spectra with a mixing time of 150 ms. NmrPipe ^57^ was used for data processing and analysis was done with Sparky ^58^. Two-dimensional {^1^H}-^15^N steady state heteronuclear NOE experiments were acquired with a 5 s relaxation delay between experiments. Standard deviations in heteronuclear NOEs were estimated based on the background noise level. Chemical shift perturbations between B_1_ and S_b1_ were calculated using Δδ_total_=((*W*_H_Δδ_H_)^2^ + (*W*_N_ΔδN)^2^)^1/2^, where *W*_H_ is 1, *W*_N_ is 0.2, and Δδ_H_ and Δδ_N_ represent ^1^H and ^15^N chemical shift changes, respectively. For PRE experiments on S_b1_, single-site cysteine mutant samples were incubated with 10 equivalents of MTSL ((1-oxyl-2,2,5,5-tetramethylpyrroline-3-methyl) methanethiosulfonate, Santa Cruz Biotechnology) at 25°C for 1 hour and completion of labeling was confirmed by MALDI mass spectrometry. Control samples were reduced with 10 equivalents of sodium ascorbate. Backbone amide peak intensities of the oxidized and reduced states were analyzed using Sparky. Three-dimensional structures were calculated with CS-Rosetta3.2 using experimental backbone ^15^N, ^1^H_N_, ^1^Hα ^13^Cα, ^13^Cβ, and ^13^CO chemical shift restraints and were either validated by comparison with experimental backbone NOE patterns (A_1_, B_1_, B_4_, S_b1_) or directly employed interproton NOEs (S_a1_, S_b2_) or PREs (S_b1_) as additional restraints. One thousand CS-Rosetta structures were calculated from which the 10 lowest energy structures were chosen. For S_b3_, CS-Rosetta failed to converge to a unique low energy topology, producing an approximately even mixture of S- and B-type folds despite the chemical shifts and NOE pattern indicating an S-fold. In this case, CNS1.1 ^59^ was employed to determine the structure as described previously ^46^, including backbone dihedral restraints from chemical shift data using TALOS ^60^. Protein structures were displayed and analyzed utilizing PROCHECK-NMR ^61^, MOLMOL ^62^ and PyMol (Schrodinger) ^45^.

## Data availability statement

The NMR structures are available in the PDB (Accession Codes: 7MN1, 7MQ4, 7MN2, 7MP7) and PDBDEV (00000083, 00000084, 00000085). All other data are available from the corresponding authors on reasonable request.

## Author contributions

B.R., Y.H., and Yw.C. contributed equally. Protein design: Yw.C., B.R., E.C., J.O., P.B.; Performed thermodynamic and binding analyses: B.R., Yw.C., R.S., P.B.; Performed dynamic light scattering experiments: T.G.; Performed NMR experiments/structural analysis: Y.H., Yh.C., T.S., T.K., J.O.; Wrote paper: J.O. (NMR and structural analysis), Yw.C., B.R., P.B. (remaining sections).

## Acknowledgments

This work was supported by National Institutes of Health Grant GM62154 (to PB and JO) and 5R44GM126676 (to PB). The NMR facility is supported by the University of Maryland, the National Institute of Standards and Technology, and a grant from the W. M. Keck Foundation. We also thank Drs. Nese Sari and Louisa P. Wu for critically reading the manuscript and many thoughtful comments. Mention of commercial products does not imply recommendation or endorsement by NIST.

## References

1 Ambroggio, X. I. & Kuhlman, B. Design of protein conformational switches. Curr Opin Struct Biol 16, 525–530 (2006).

2 Bryan, P. N. & Orban, J. Proteins that switch folds. Curr. Opin. Struct. Biol. 20, 482–488 (2010).

3 Dishman, A. F. et al. Evolution of fold switching in a metamorphic protein. Science 371, 86–90, doi:10.1126/science.abd8700 (2021).

4 Wei, K. Y. et al. Computational design of closely related proteins that adopt two well-defined but structurally divergent folds. Proc Natl Acad Sci U S A 117, 7208–7215, doi:10.1073/pnas.1914808117 (2020).

5 Anderson, W. J., Van Dorn, L. O., Ingram, W. M. & Cordes, M. H. Evolutionary bridges to new protein folds: design of C-terminal Cro protein chameleon sequences. Protein Eng Des Sel 24, 765–771, doi:10.1093/protein/gzr027 (2011).

6 Burmann, B. M. et al. An α helix to β barrel domain switch transforms the transcription factor RfaH into a translation factor. Cell 150, 291–303, doi:10.1016/j.cell.2012.05.042 (2012).

7 Kulkarni, P. et al. Structural metamorphism and polymorphism in proteins on the brink of thermodynamic stability. Protein Sci 27, 1557–1567, doi:10.1002/pro.3458 (2018).

8 Dishman, A. F. & Volkman, B. F. Design and discovery of metamorphic proteins. Curr Opin Struct Biol 74, 102380, doi:10.1016/j.sbi.2022.102380 (2022).

9 Rackovsky, S. Nonlinearities in protein space limit the utility of informatics in protein biophysics. Proteins 83, 1923–1928, doi:10.1002/prot.24916 (2015).

10 Chen, S. H., Meller, J. & Elber, R. Comprehensive analysis of sequences of a protein switch. Protein Sci 25, 135–146, doi:10.1002/pro.2723 (2016).

11 Li, W., Kinch, L. N., Karplus, P. A. & Grishin, N. V. ChSeq: A database of chameleon sequences. Protein Sci 24, 1075–1086, doi:10.1002/pro.2689 (2015).

12 Wolynes, P. G. Evolution, energy landscapes and the paradoxes of protein folding. Biochimie 119, 218–230, doi:10.1016/j.biochi.2014.12.007 (2015).

13 Holzgräfe, C. & Wallin, S. Smooth functional transition along a mutational pathway with an abrupt protein fold switch. Biophys J 107, 1217–1225, doi:10.1016/j.bpj.2014.07.020 (2014).

14 Scheraga, H. A. & Rackovsky, S. Homolog detection using global sequence properties suggests an alternate view of structural encoding in protein sequences. Proc Natl Acad Sci U S A 111, 5225–5229, doi:10.1073/pnas.1403599111 (2014).

15 Ha, J. H. & Loh, S. N. Protein conformational switches: from nature to design. Chemistry 18, 7984–7999, doi:10.1002/chem.201200348 (2012).

16 Yadid, I., Kirshenbaum, N., Sharon, M., Dym, O. & Tawfik, D. S. Metamorphic proteins mediate evolutionary transitions of structure. Proc Natl Acad Sci U S A 107, 7287–7292, doi:10.1073/pnas.0912616107 (2010).

17 Lichtarge, O. & Wilkins, A. Evolution: a guide to perturb protein function and networks. Curr Opin Struct Biol 20, 351–359, doi:10.1016/j.sbi.2010.04.002 (2010).

18 Rollins, N. J. et al. Inferring protein 3D structure from deep mutation scans. Nat Genet 51, 1170–1176, doi:10.1038/s41588-019-0432-9 (2019).

19 Sikosek, T., Chan, H. S. & Bornberg-Bauer, E. Escape from Adaptive Conflict follows from weak functional trade-offs and mutational robustness. Proc Natl Acad Sci U S A 109, 14888–14893, doi:10.1073/pnas.1115620109 (2012).

20 Chen, N., Das, M., LiWang, A. & Wang, L. P. Sequence-Based Prediction of Metamorphic Behavior in Proteins. Biophys J 119, 1380–1390, doi:10.1016/j.bpj.2020.07.034 (2020).

21 Porter, L. L. & Looger, L. L. Extant fold-switching proteins are widespread. Proc Natl Acad Sci U S A 115, 5968–5973, doi:10.1073/pnas.1800168115 (2018).

22 Bedford, J. T., Poutsma, J., Diawara, N. & Greene, L. H. The nature of persistent interactions in two model β-grasp proteins reveals the advantage of symmetry in stability. Journal of Computational Chemistry 42, 600–607, doi:https://doi.org/10.1002/jcc.26477 (2021).

23 Falkenberg, C., Bjorck, L. & Akerstrom, B. Localization of the binding site for streptococcal protein G on human serum albumin. Identification of a 5.5-kilodalton protein G binding albumin fragment. Biochemistry 31, 1451–1457 (1992).

24 Frick, I. M. et al. Convergent evolution among immunoglobulin G-binding bacterial proteins. Proc Natl Acad Sci U S A 89, 8532–8536 (1992).

25 Myhre, E. B. & Kronvall, G. Heterogeneity of nonimmune immunoglobulin Fc reactivity among gram-positive cocci: description of three major types of receptors for human immunoglobulin G. Infect. Immun. 17, 475–482 (1977).

26 Reis, K. J., Ayoub, E. M. & Boyle, M. D. P. Streptococcal Fc receptors. II. Comparison of the reactivity of a receptor from a group C streptococcus with staphylococcal protein A. J. Immunol. 132, 3098–3102 (1984).

27 Lindberg, M. O., Haglund, E., Hubner, I. A., Shakhnovich, E. I. & Oliveberg, M. Identification of the minimal protein-folding nucleus through loop-entropy perturbations. Proc Natl Acad Sci U S A 103, 4083–4088, doi:10.1073/pnas.0508863103 (2006).

28 Haglund, E., Lindberg, M. O. & Oliveberg, M. Changes of protein folding pathways by circular permutation. Overlapping nuclei promote global cooperativity. J Biol Chem 283, 27904–27915, doi:10.1074/jbc.M801776200 (2008).

29 Haglund, E. et al. The HD-exchange motions of ribosomal protein S6 are insensitive to reversal of the protein-folding pathway. Proc Natl Acad Sci U S A 106, 21619–21624, doi:10.1073/pnas.0907665106 (2009).

30 Haglund, E. et al. Trimming down a protein structure to its bare foldons: spatial organization of the cooperative unit. J Biol Chem 287, 2731–2738, doi:10.1074/jbc.M111.312447 (2012).

31 Lindahl, M. et al. Crystal structure of the ribosomal protein S6 from Thermus thermophilus. Embo j 13, 1249–1254, doi:10.2210/pdb1ris/pdb (1994).

32 Day, R., Beck, D. A., Armen, R. S. & Daggett, V. A consensus view of fold space: combining SCOP, CATH, and the Dali Domain Dictionary. Protein Sci 12, 2150–2160, doi:10.1110/ps.0306803 (2003).

33 He, Y., Chen, Y., Alexander, P., Bryan, P. N. & Orban, J. NMR structures of two designed proteins with high sequence identity but different fold and function. Proc Natl Acad Sci U S A 105, 14412–14417 (2008).

34 Alexander, P. A., He, Y., Chen, Y., Orban, J. & Bryan, P. N. A minimal sequence code for switching protein structure and function. Proc Natl Acad Sci U S A 106, 21149–21154 (2009).

35 He, Y., Chen, Y., Alexander, P. A., Bryan, P. N. & Orban, J. Mutational tipping points for switching protein folds and functions. Structure 20, 283–291 (2012).

36 Gallagher, T. D., Gilliland, G., Wang, L. & Bryan, P. The prosegment-subtilisin BPN’ complex: crystal structure of a specific foldase. Structure 3, 907–914 (1995).

37 Tangrea, M. A. et al. Stability and global fold of the mouse prohormone convertase 1 pro-domain. Biochemistry 40, 5488–5495. (2001).

38 Tangrea, M. A., Bryan, P. N., Sari, N. & Orban, J. Solution Structure of the Pro-hormone Convertase 1 Pro-domain from Mus musculus. J Mol Biol 320, 801–812 (2002).

39 Sari, N. et al. Hydrogen-deuterium exchange in free and prodomain-complexed subtilisin. Biochemistry 46, 652–658 (2007).

40 Orengo, C. A. & Thornton, J. M. Alpha plus beta folds revisited: some favoured motifs. Structure 1, 105–120, doi:10.1016/0969-2126(93)90026-d (1993).

41 Chen, Y. et al. Engineering subtilisin proteases that specifically degrade active RAS. Communications Biology 4, 299, doi:10.1038/s42003-021-01818-7 (2021).

42 Alexander, P. A., Rozak, D. A., Orban, J. & Bryan, P. N. Directed evolution of highly homologous proteins with different folds by phage display: implications for the protein folding code. Biochemistry 44, 14045–14054 (2005).

43 Alexander, P. A., He, Y., Chen, Y., Orban, J. & Bryan, P. N. The design and characterization of two proteins with 88% sequence identity but different structure and function. Proc Natl Acad Sci U S A 104, 11963–11968 (2007).

44 Leaver-Fay, A. et al. ROSETTA3: an object-oriented software suite for the simulation and design of macromolecules. Methods Enzymol 487, 545–574 (2011).

45 The PyMOL Molecular Graphics System (DeLano Scientific, San Carlos, CA, 2002).

46 He, Y. et al. Structure, dynamics, and stability variation in bacterial albumin binding modules: implications for species specificity. Biochemistry 45, 10102–10109 (2006).

47 Shen, Y. et al. De novo structure generation using chemical shifts for proteins with high-sequence identity but different folds. Protein Sci. 19, 349–356 (2010).

48 Chen, Y. et al. Rules for designing protein fold switches and their implications for the folding code. bioRxiv, 2021.2005.2018.444643, doi:10.1101/2021.05.18.444643 (2021).

49 Rozak, D. A., Orban, J. & Bryan, P. N. G148-GA3: a streptococcal virulence module with atypical thermodynamics of folding optimally binds human serum albumin at physiological temperatures. Biochim Biophys Acta 1753, 226–233 (2005).

50 He, Y., Chen, Y., Rozak, D. A., Bryan, P. N. & Orban, J. An artificially evolved albumin binding module facilitates chemical shift epitope mapping of GA domain interactions with phylogenetically diverse albumins. Protein Sci 16, 1490–1494 (2007).

51 He, Y. et al. Solution NMR structure of a sheddase inhibitor prodomain from the malarial parasite Plasmodium falciparum. Proteins 80, 2810–2817, doi:10.1002/prot.24187 (2012).

52 Alexander, P., Fahnestock, S., Lee, T., Orban, J. & Bryan, P. Thermodynamic analysis of the folding of the Streptococcal Protein G IgG-binding domains B1 and B2: why small proteins tend to have high denaturation temperatures. Biochemistry 31, 3597–3603 (1992).

53 Chien, S.-C., Chen, C.-Y., Lin, C.-F. & Yeh, H.-I. Critical appraisal of the role of serum albumin in cardiovascular disease. Biomarker Research 5, 31, doi:10.1186/s40364-017-0111-x (2017).

54 Gonzalez-Quintela, A. et al. Serum levels of immunoglobulins (IgG, IgA, IgM) in a general adult population and their relationship with alcohol consumption, smoking and common metabolic abnormalities. Clin Exp Immunol 151, 42–50, doi:10.1111/j.1365-2249.2007.03545.x (2008).

55 Studier, F. W. Protein production by auto-induction in high density shaking cultures. Protein Expr Purif 41, 207–234 (2005).

56 Ruan, B., Fisher, K. E., Alexander, P. A., Doroshko, V. & Bryan, P. N. Engineering subtilisin into a fluoride-triggered processing protease useful for one-step protein purification. Biochemistry 43, 14539–14546 (2004).

57 Delaglio, F. et al. NMRPipe: a multidimensional spectral processing system based on UNIX pipes. J. Biomol. NMR 6, 277–293 (1995).

58 SPARKY 3 v. 3 (University of California San Francisco, 2004).

59 Brunger, A. T. et al. Crystallography & NMR system: A new software suite for macromolecular structure determination. Acta Crystallogr. D (Biol. Crystallogr.) 54, 905–921 (1998).

60 Cornilescu, G., Delaglio, F. & Bax, A. Protein backbone angle restraints from searching a database for chemical shift and sequence homology. J. Biomol. NMR 13, 289–302 (1999).

61 Laskowski, R. A., Rullmann, J. A., MacArthur, M. W., Kaptein, R. & Thornton, J. M. AQUA and PROCHECK-NMR: Programs for checking the quality of protein structures solved by NMR. J. Biomol. NMR 8, 477–486 (1996).

62 Koradi, R., Billeter, M. & Wuthrich, K. MOLMOL: a program for display and analysis of macromolecular structures. J Mol Graph 14, 51–55 (1996).

