## Supplement for "Design and characterization of a protein fold switching network"

**Table S1.** Fold, stability, and binding properties for each designed protein.

| Mutant | Fold | $\Delta G_{folding}$ | Binding ligand | $K_I$ and or $K_D$ |
|---|---|---|---|---|
| SI | S | -8.5 kcal/mol | Protease | 0.2μM |
| $S_{a1}I$ | S | -5.3 kcal/mol | Protease | 0.05μM |
| $S_{a1}$ | S | -5.3 kcal/mol | --- | |
| $A_1$ | A | -4 kcal/mol | --- | |
| $S_{a2}I$ | S | -5.3 kcal/mol | Protease/HSA | 0.05μM/100μM |
| $A_2$ | A | -4 kcal/mol | HSA | ≤ 1μM |
| $S_{b1}$ | B | -1.1 kcal/mol | --- | |
| $B_1$ | B | -4 kcal/mol | --- | |
| $S_{b2}$ | S | -4 kcal/mol | --- | |
| $B_2$ | unfolded | >2 kcal/mol | --- | |
| S'I | S | -8.5 kcal/mol | Protease | 0.06μM |
| $S_{b3}I$ | S | -3.5 kcal/mol | Protease/IgG | 0.05μM/10μM |
| $S_{b3}$ | S | -3.5 kcal/mol | IgG | 10μM |
| $B_3$ | B | -1.2 kcal/mol | IgG | ≤ 1μM |
| $S_{b4}$ | Mixed S/B | 0 kcal/mol | IgG | ≤ 1μM |
| $B_4$ | B | -4.1 kcal/mol | IgG | ≤ 1μM |
| $S_{b5}$ | B | -5 kcal/mol | IgG | 10μM |

$\Delta G_{folding}$ 25˚C [1]

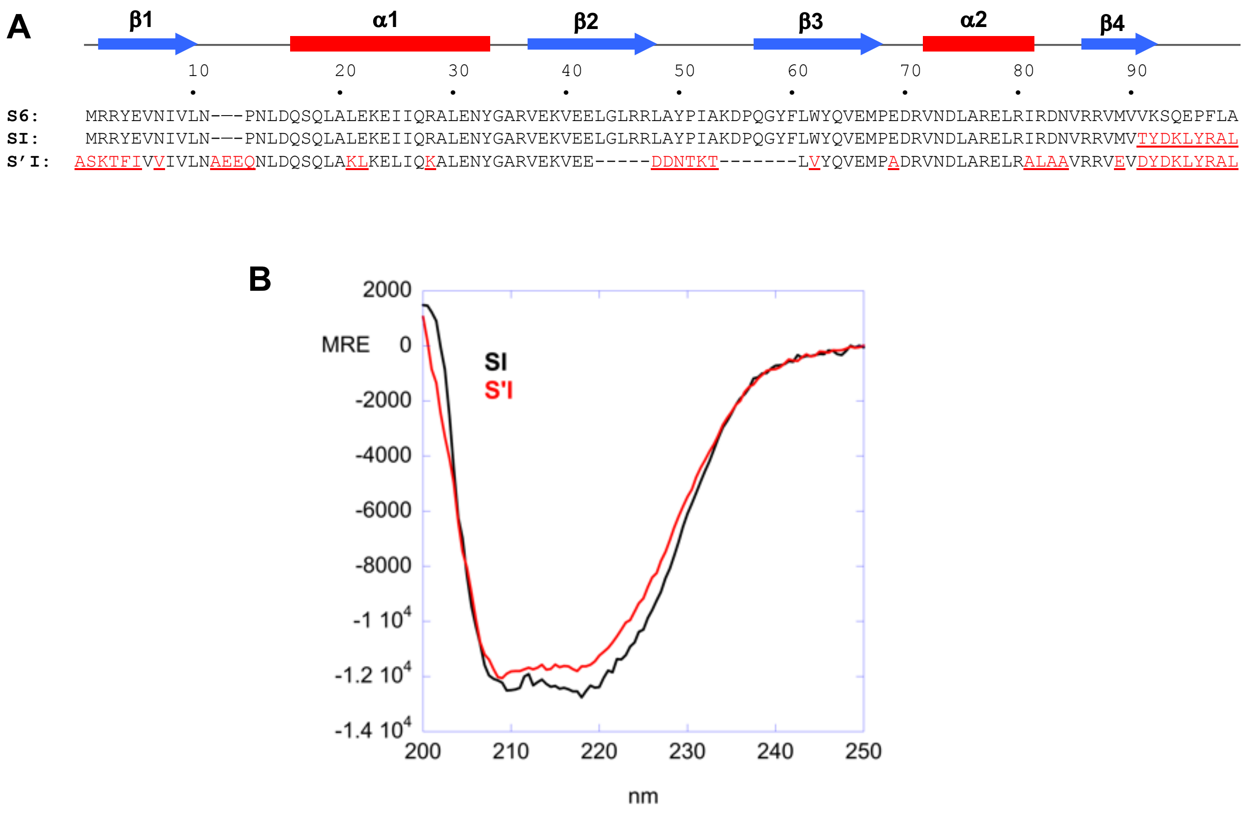


**Fig. S1:** Sequence comparison of S6, SI, and S'I. The secondary structure for the parent S6 fold is shown above the alignment. Mutations are shown by red text. CD spectra are compared in panel B.

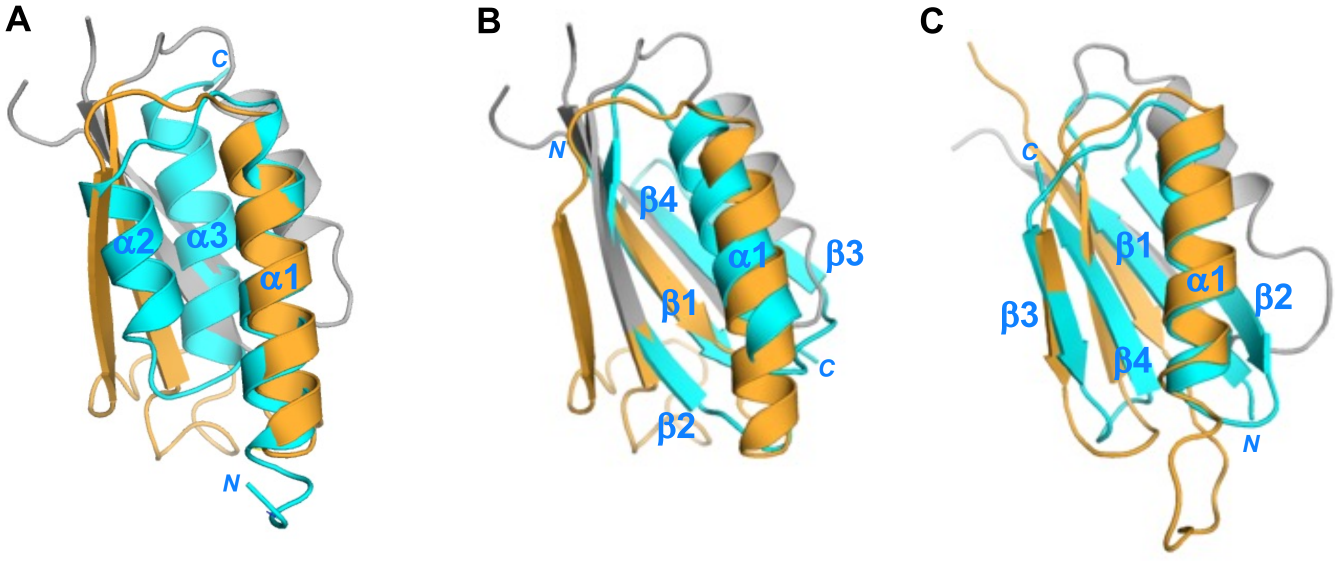


**S/A design**
*(α1 aligned)*

**S/B design**
*(β1 aligned)*

**S'/B design**
*(α1/β3/β4 aligned)*

**Fig. S2.** Topological alignment of A and B-folds with S-folds. (A) The embedded A-fold occupies amino acid positions 11-66 with 46% identity in the aligned regions for the parent sequences. α1(S) mostly coincides with α1(A). β2(S) becomes α2(A), and the long β2-β3 turn and first half of β3(S) becomes α3(A). β1 and α2-β4 of the S-fold are outside of the overlap region. (B) The embedded B-fold occupies amino acid positions 4-59 with 9% identity in the aligned regions for the parent sequences. β1(S) mostly coincides with β1(B). The first half of α1(S) becomes β2(B) and the second half of α1(S) becomes the first half of α1(B). The β2 strand of S becomes the second half of α1(B), a turn, and the first part of β3(B). The long β2-β3(S) turn and the first part of β3(S) become the second part of β3 and β4 of B. The second half of β3 and α2-β4 of the S-fold are outside of the overlap region. (C) The embedded B-fold occupies amino acid positions 1-56 with 16% identity in the aligned regions for the parent sequences (excluding the 3 residue insertion and 12 residue deletion). The β1–strand is the same in both folds but changes orientation, the long turn between β1 and α1 in $S_{b3}$ becomes β2 of the B-fold, and the α1–β2–β3 of $S_{b3}$ maintains the same topology in both folds. The α2-helix and the β4–strand of $S_{b3}$ are outside the overlap region.

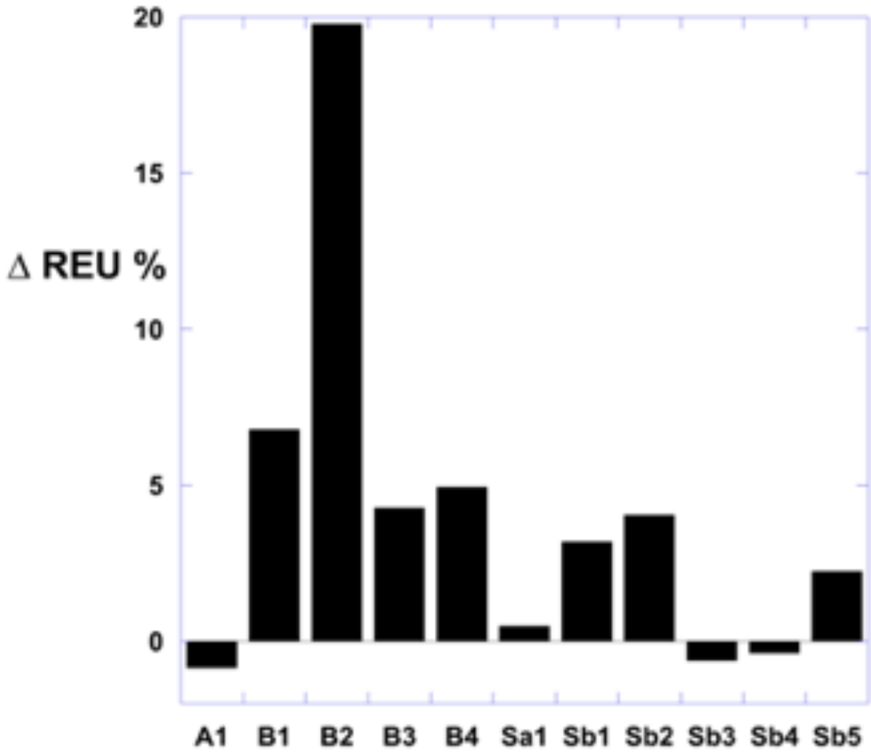


**Fig. S3:** The percent change in Rosetta Energy Units (REU) between the parent protein and the designed switch protein is plotted. The computational design of $A_1$ was compared to the relaxed structure of a highly stable A-fold [2]. The designs of $B_1$, $B_2$, $B_3$, and $B_4$ were compared to a highly stable B-fold [3]. The designs of $S_{a1}$, $S_{b1}$, $S_{b2}$, $S_{b3}$, $S_{b4}$, and $S_{b5}$ were compared to a highly stable S-fold [4]. All designed proteins have relatively small changes in %ΔREU except for $B_2$. The 20% increase in REU for $B_2$ is consistent with its low stability.

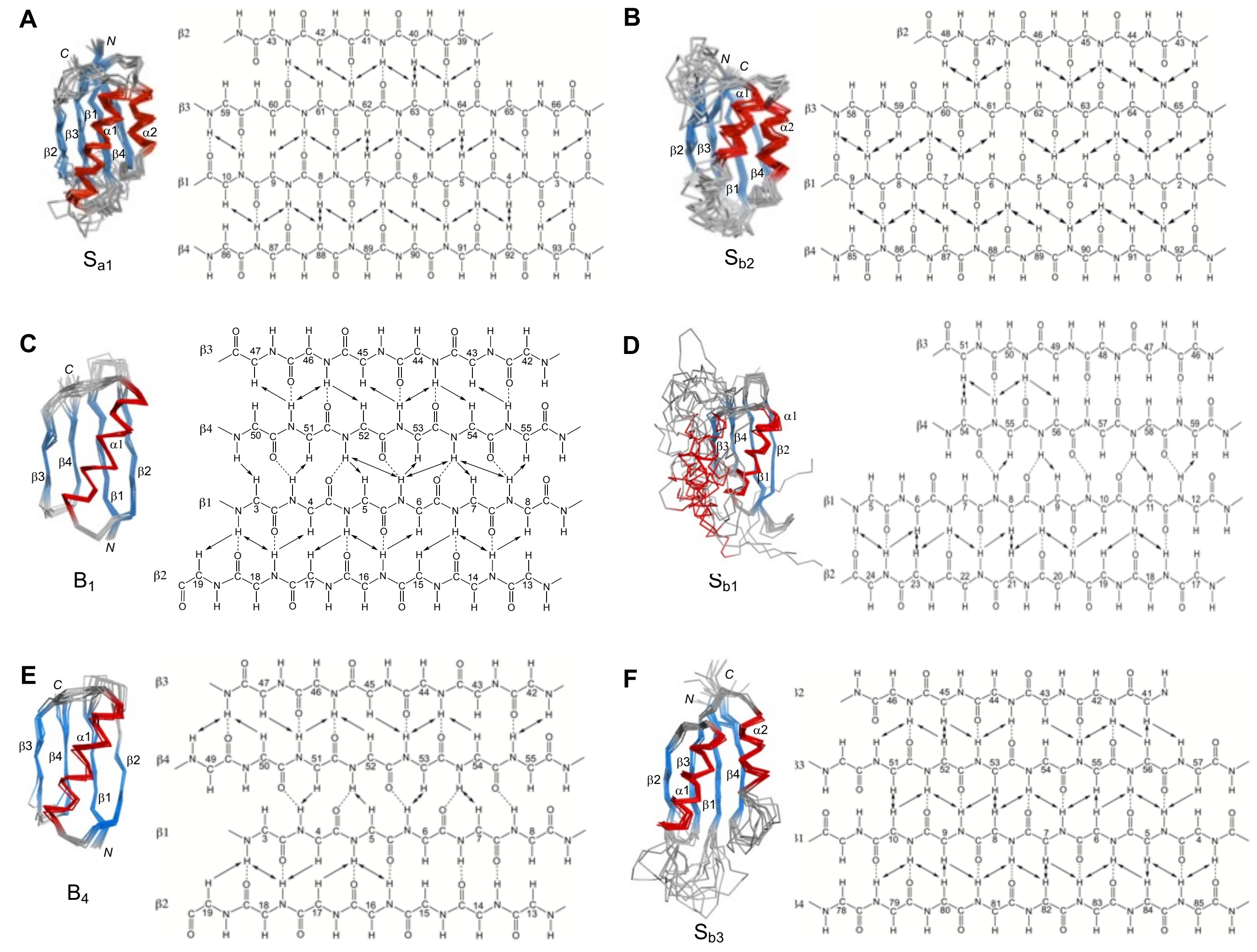


**Fig. S4:** Summary of long-range backbone NOEs observed for β-sheets in designed proteins**.**

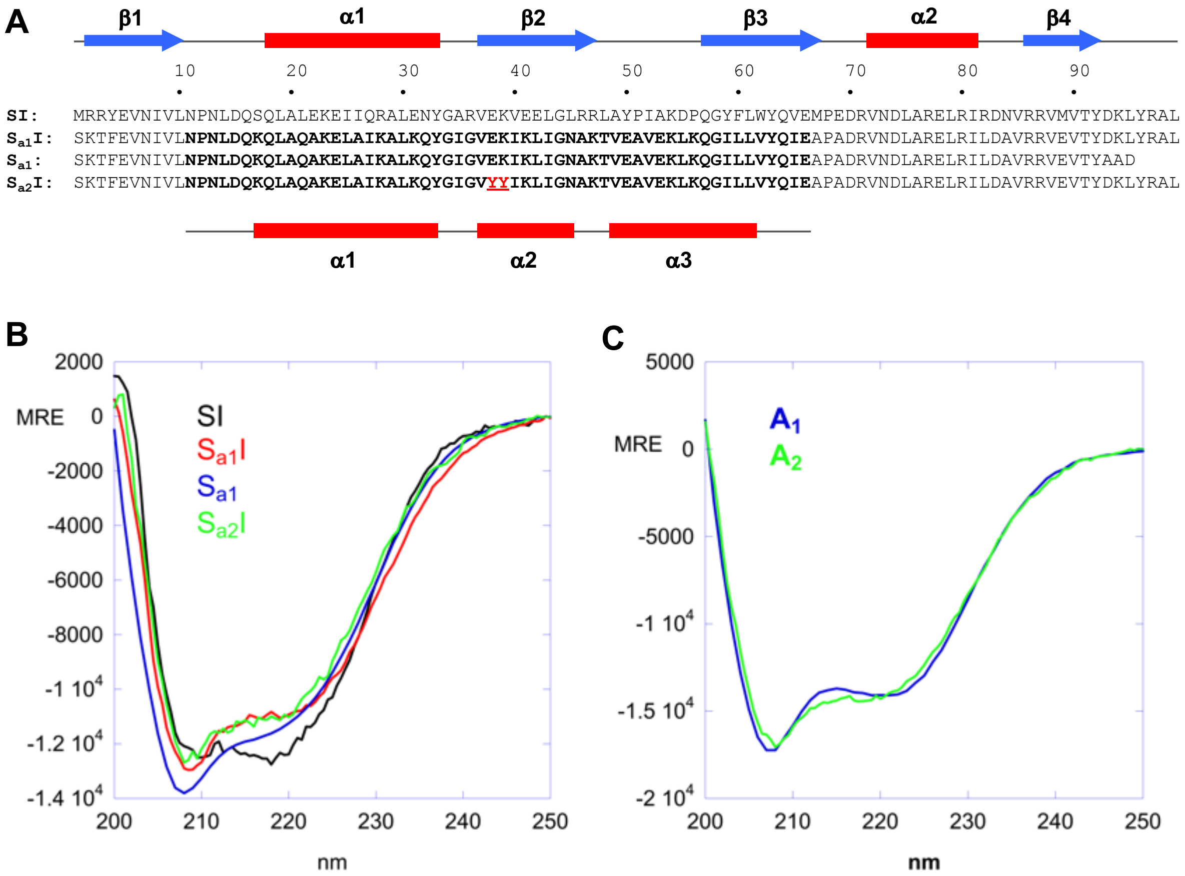


**Fig S5:** Sequence comparison of SI, $S_{a1}$, $S_{a2}$, and $S_{a2}$I. The secondary structures for the parent S6 and A folds are shown above and below the alignment, respectively. Mutations are shown by red text. CD spectra are compared in panels B and C.

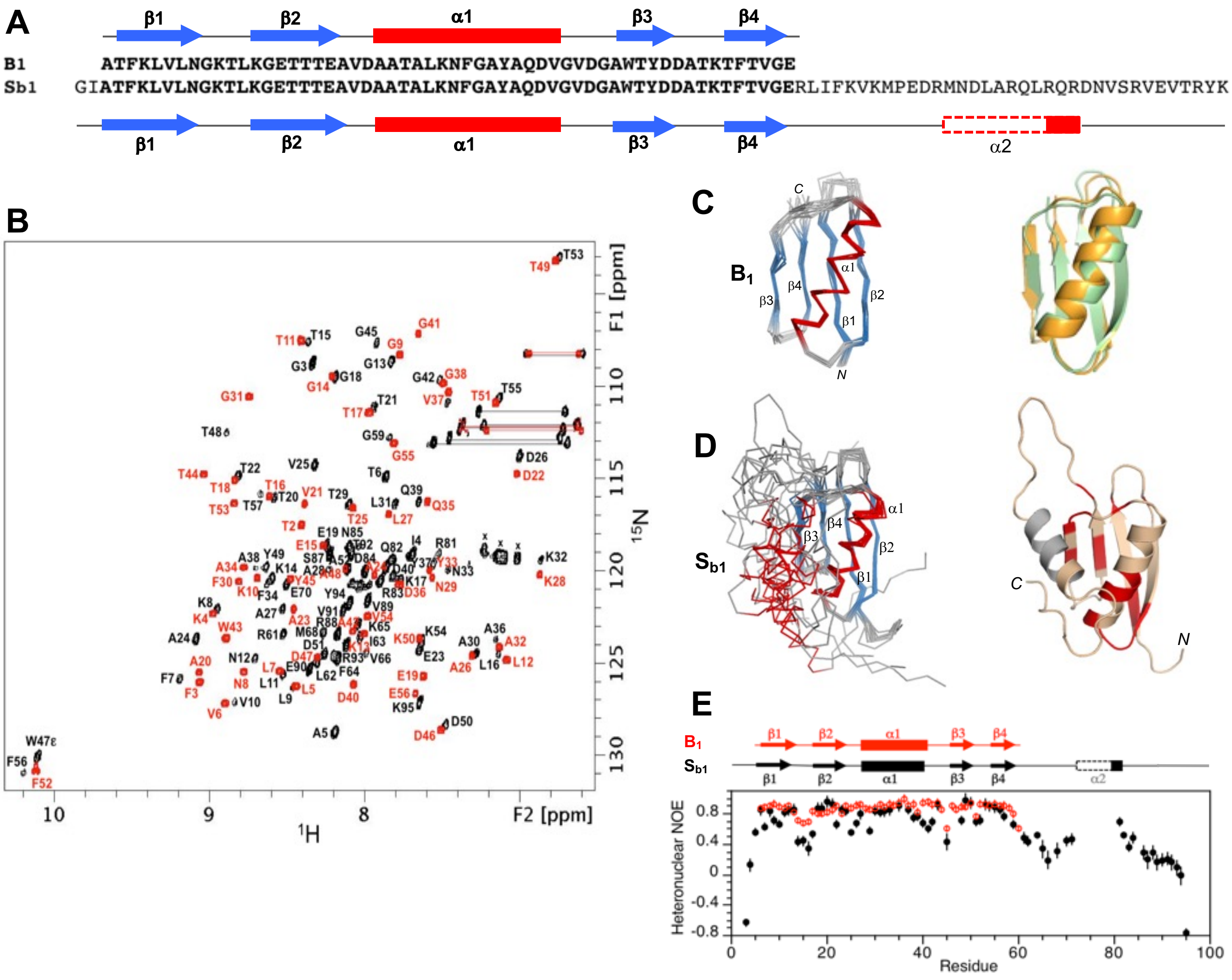


**Figure S6**: Structure and dynamics of $B_1$ and $S_{b1}$ - $S_{b1}$ has a B-fold. (A) Sequence alignment of $B_1$ and $S_{b1}$, which are 100% identical over the B-region. (B) Overlaid two dimensional $^{1}H$-$^{15}N$ HSQC spectra of $S_{b1}$ (black) and $B_1$ (red) with backbone amide assignments. Spectra were recorded at 10ºC. (C) Ensemble of 10 lowest energy CS-Rosetta structures for $B_1$ (left panel). Superposition of the $B_1$ structure (green) with the parent GB1 fold (orange) (right panel). (D) Ensemble of 10 lowest energy CS-Rosetta structures for $S_{b1}$ using chemical shift, NOE, and PRE restraints (left panel). Cartoon representation of model 1 from the ensemble (right panel). Values of $\Delta\delta_{total}$>0.1 ppm from Fig. S7A are mapped onto the structure (red). Unassigned residues in the putative $\alpha$2-helix are in gray. (E) Plot of {$^{1}H$}-$^{15}N$ steady state heteronuclear NOE values at 600 MHz versus residue for $B_1$ (red) and for $S_{b1}$ (black). Error bars indicate ±1SD.

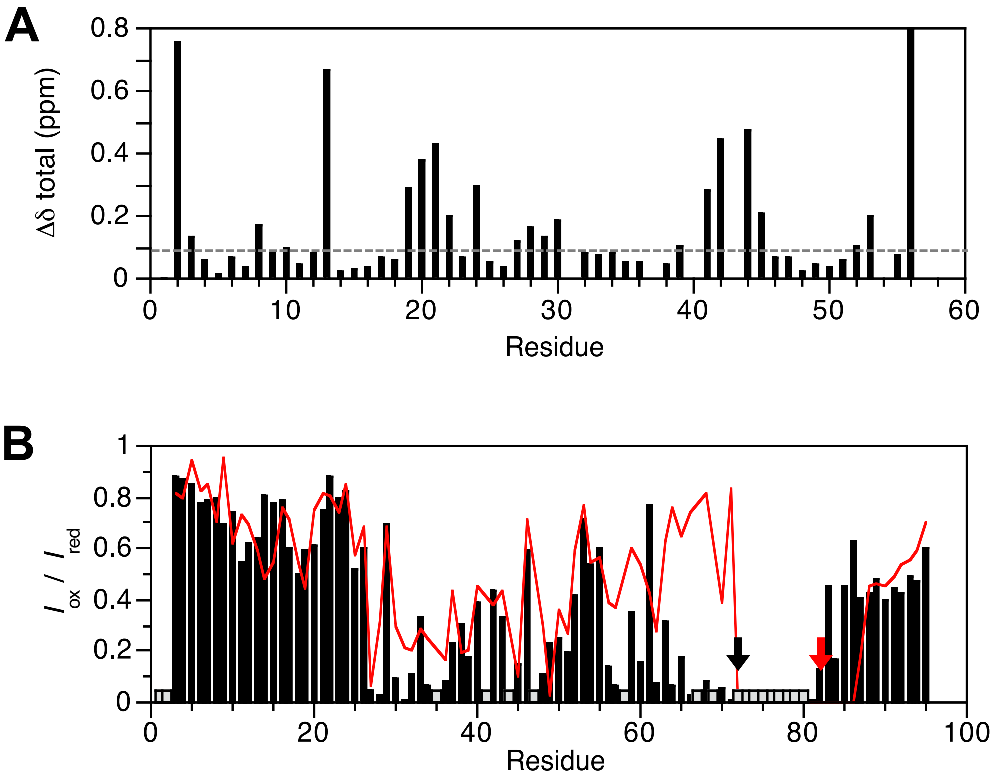


**Figure S7**: Chemical shift perturbation (CSP) and PRE profiles for $S_{b1}$. (A) Plot of CSPs between $S_{b1}$ and $B_1$ for backbone amides in the 56 amino acid identical region. Residue numbering is for $B_1$. (B) Plot of $I_{ox}/I_{red}$ versus residue for $S_{b1}$-R72C-MTSL (black) and $S_{b1}$-R83C-MTSL (red). Similar PRE profiles are observed in the B-folded part of the structure when the spin label is attached to the N- or C-terminus of the exchange-broadened region, which corresponds with $\alpha 2$ in S6. Gray columns indicate unassigned residues or prolines. The positions of the spin labels are indicated with arrows.

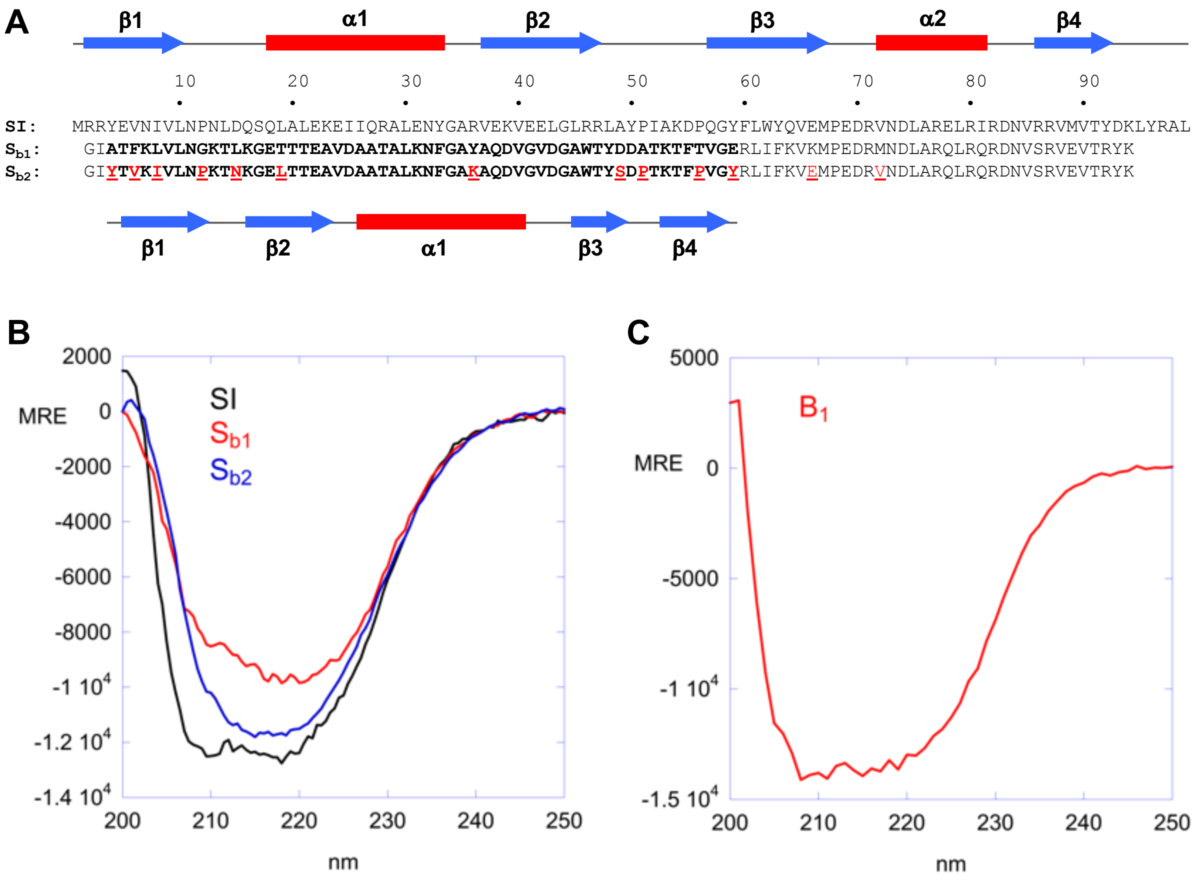


**Fig. S8**: Sequence comparison of SI, $S_{b1}$, and $S_{b2}$. The secondary structures for the parent S6 and B folds are shown above and below the alignment, respectively. Mutations are shown by red text. CD spectra are compared in panels B and C.

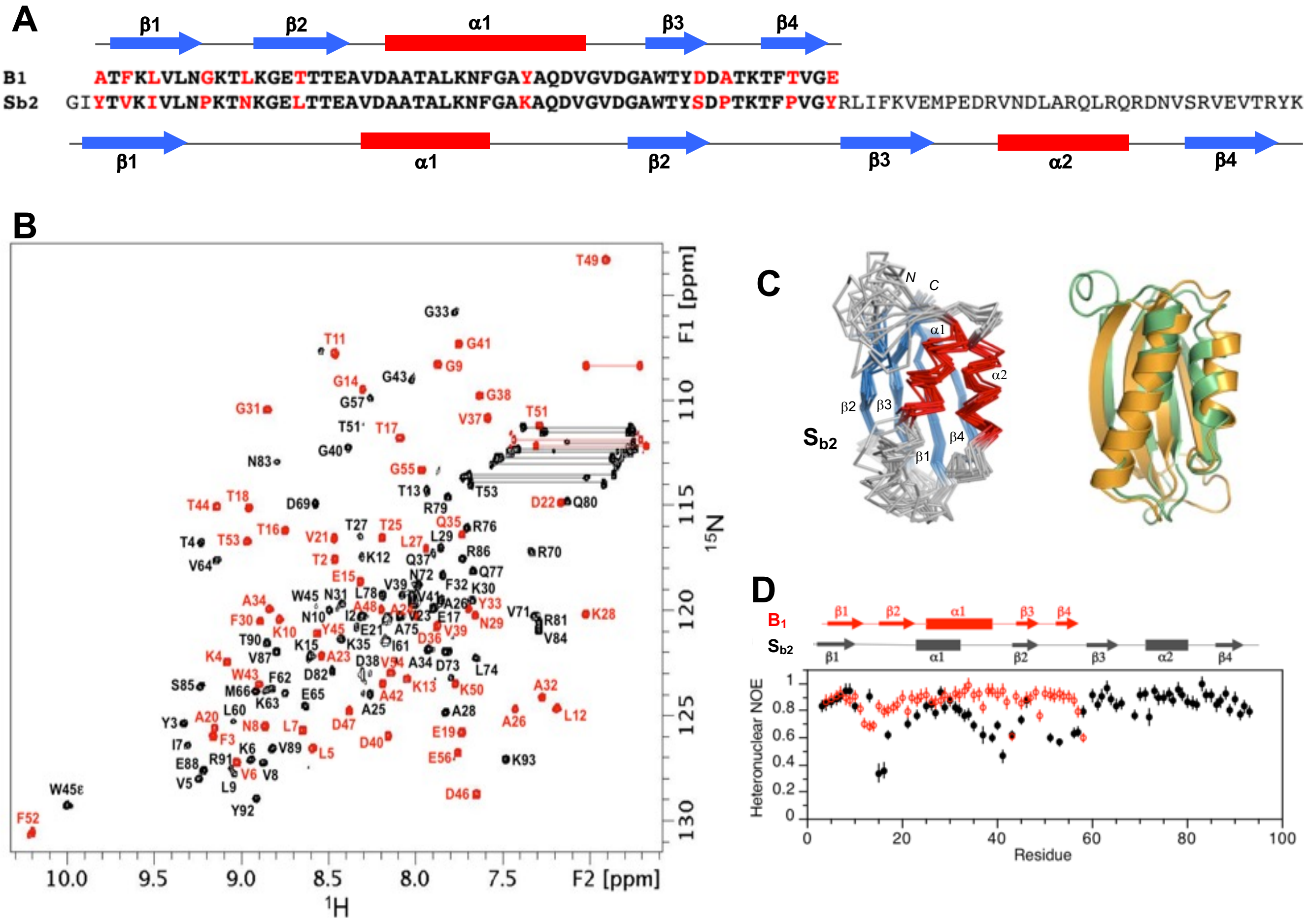


**Figure S9**: Structure and dynamics of $S_{b2}$ – $S_{b2}$ has an S-fold. (A) Sequence alignment of $S_{b2}$ and $B_1$. The 11 amino acid differences are shown in red. (B) Overlaid two dimensional $^1H$-$^{15}N$ HSQC spectra of $S_{b2}$ (black) and $B_1$ (red) with backbone amide assignments. Spectra were recorded at 25°C. (C) Ensemble of 10 lowest energy CS-Rosetta structures for $S_{b2}$ (left panel). Superposition of $S_{b2}$ (green) with the parent S6 fold (orange) (right panel). (D) Plot of {$^1H$}-$^{15}N$ steady state heteronuclear NOE values at 600 MHz versus residue for $S_{b2}$ (black). Values are compared with $B_1$, which has 80% sequence identity in the corresponding region. Error bars indicate ±1SD.

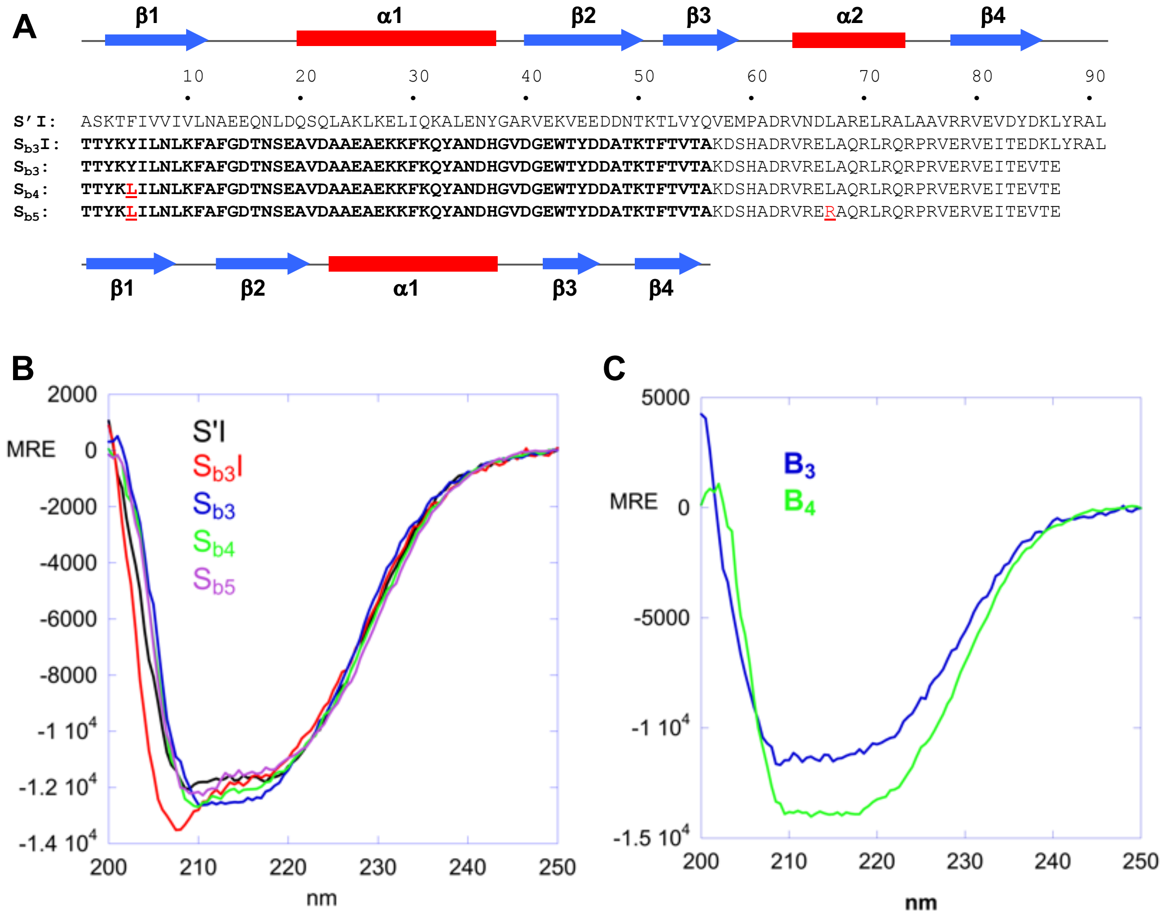


**Fig. S10**: Sequence comparison of S'I, $S_{b3}$, $S_{b4}$, and $S_{b5}$. The secondary structures for the parent S6 and B folds are shown above and below the alignment, respectively. Mutations are shown by red text. CD spectra are compared in panels B and C.

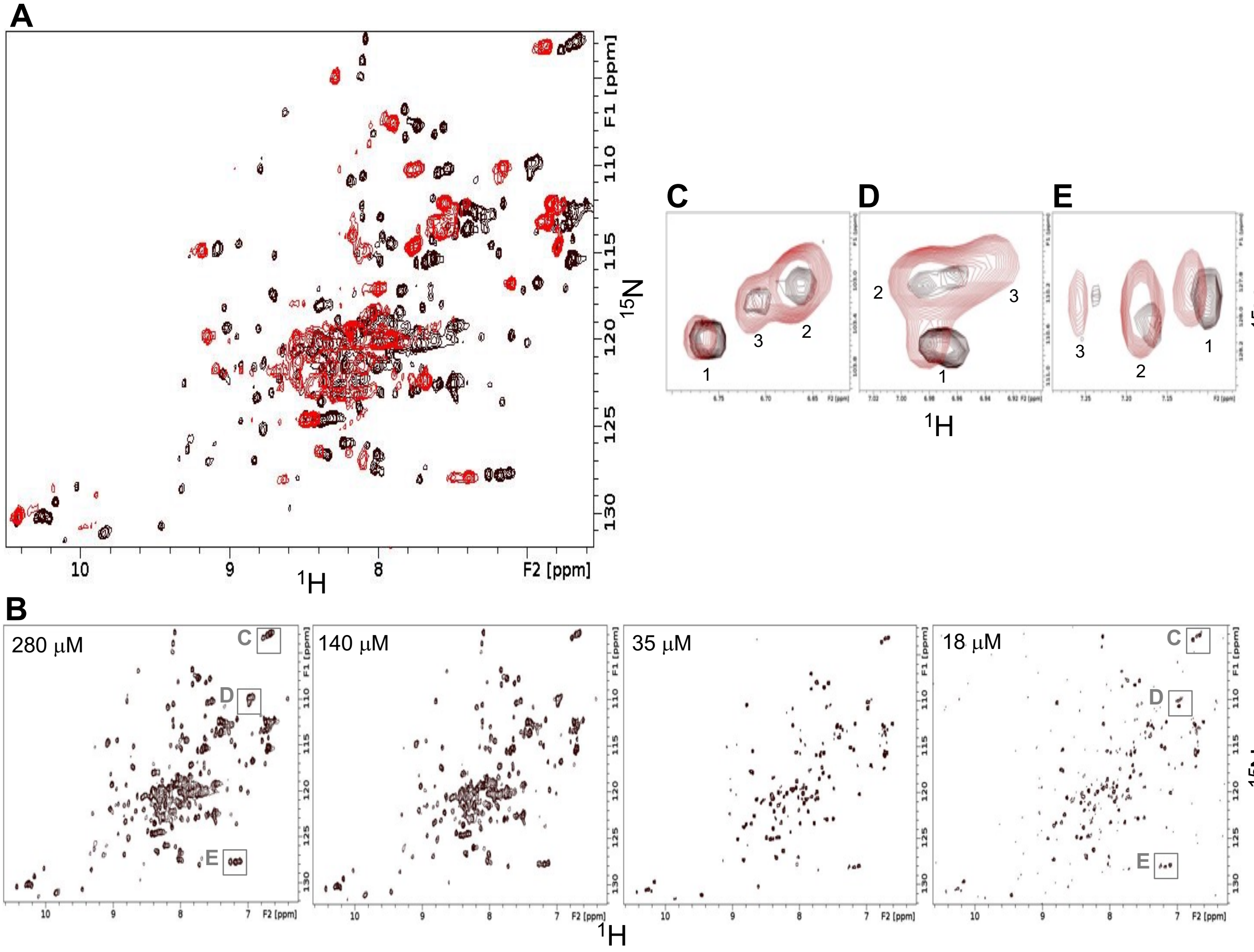


**Fig. S11:** Temperature and concentration dependence of $B_3$. (A) Overlaid two dimensional $^1H$-$^{15}N$ HSQC spectra of $B_3$ at 25°C (red) and at 5°C (black). The protein concentration in both spectra is 280 μM. Broadened peaks at 25°C are consistent with low stability. (B) HSQC spectra of $B_3$ recorded at different concentrations as indicated. All spectra were acquired at 5°C. (C, D, E) Expanded regions from the overlaid concentration dependent spectra of $B_3$ at 18 μM (black) and 280 μM (red). Regions are as indicated in the leftmost and rightmost panels in (B). The peaks in (C, D, E) are labeled 1, 2 and 3, corresponding to three main concentration-dependent species. At 18 μM, the putatively monomeric species 1 is dominant, while at 280 μM the putatively dimeric species 2 is dominant.

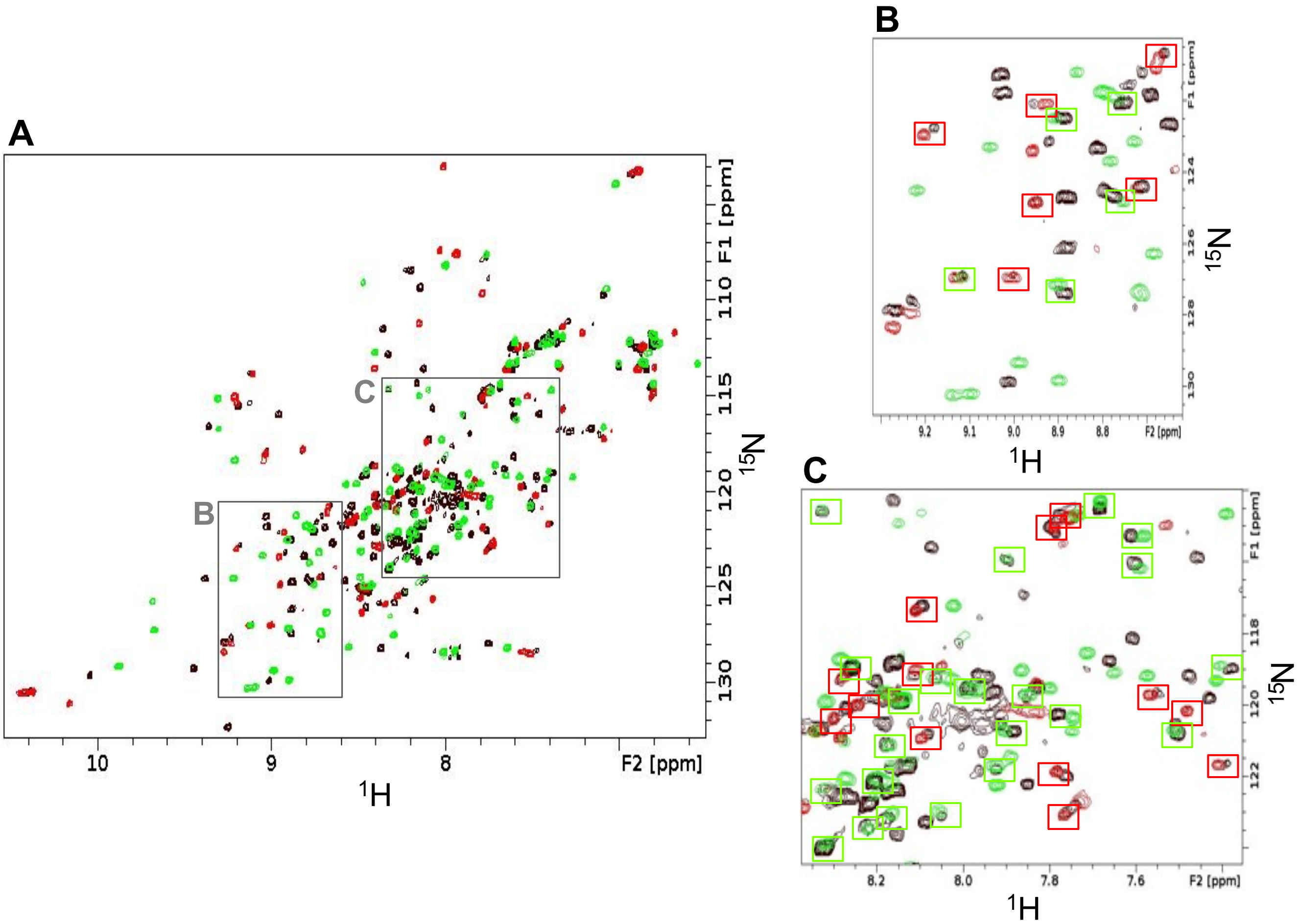


**Fig. S12:** Comparison of spectra for $S_{b4}$, $B_4$, and $S_{b3}$. (A) Two dimensional $^{1}H$-$^{15}N$ HSQC spectrum of $S_{b4}$ (black) overlaid with spectra for $B_4$ (red) and $S_{b3}$ (green). All spectra were acquired at 25ºC. (B, C) Expanded regions as indicated in (A). Overlapping or proximal $B_4$/$S_{b4}$ peaks (red boxes) and $S_{b3}$/$S_{b4}$ peaks (green boxes) are highlighted. The overlap of the spectra indicates that $S_{b4}$ populates both the S- and B-folds simultaneously. About 50 peaks in $S_{b4}$ superimpose well with $B_4$ although some peaks are shifted slightly, presumably due to the presence of a disordered C-terminal tail in $S_{b4}$ when the polypeptide chain adopts the B-fold. A significant number of $S_{b4}$ signals also coincide approximately with the $S_{b3}$ spectrum. Differences in peak positions between $S_{b4}$ and $S_{b3}$ are likely due to the L5Y mutation, which affects numerous neighboring residues.

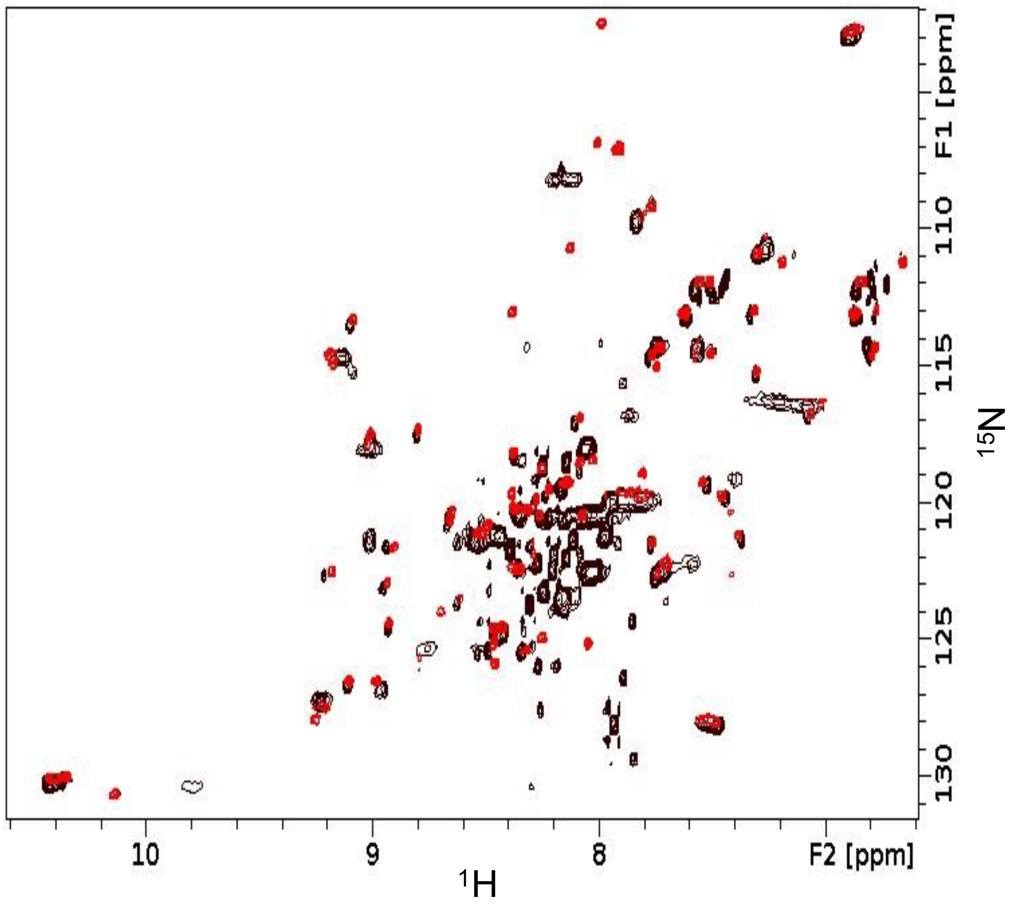


**Fig. S13**: Two dimensional $^{1}H$-$^{15}N$ HSQC spectrum of $S_{b5}$ (black) compared with $B_4$ (red). Both spectra were acquired at 25°C. The significant overlap of folded signals in the two spectra indicates that $S_{b5}$ has a B-fold with a disordered C-terminal tail.

**Structure of** $B_1$**.** The NMR structure consists of four β-strands defined by residues 2-9 (β1), residues 13-20 (β2), residues 42-46 (β3), and residues 50-55 (β4) and one α-helix from residues 23-37 (**Fig. S6B,C, Table 1**). The NOE pattern between main chain protons agrees well with a B-fold (**Fig. S4C).** Steady-state $\{^1H\}$-$^{15}N$ heteronuclear NOE values for $B_1$ are consistent with a well ordered structure, with the exception of the β1-β2, α1-β3, and β3-β4 internal loops (**Fig. S6E**).

**Structure of** $S_{b1}$**.** The topology of $S_{b1}$ is not the same as the parent S6 structure. Instead, the 2D $^1H$-$^{15}N$ HSQC spectrum of $S_{b1}$ has a pattern similar to that of $B_1$ (**Fig. S6B)**. NMR assignment of the main chain resonances showed the presence of four β-strands and two α-helices, but the order of the secondary structure elements was ββαββα rather than the βαββαβ arrangement expected for an S-type fold. Initial NMR structures of $S_{b1}$ indicated a B-fold, which was supported by backbone NOE connectivities (**Fig. S4D**), with a mostly disordered C-terminal tail. CS-Rosetta modeled residues 73-83 near the C-terminus as an α2-helix. Of these, amide signals due to residues 73-80 were not detectable in NMR spectra while residues 81-83 were helical based on assigned chemical shifts. Comparison of $S_{b1}$ amide chemical shifts with those of $B_1$ indicated that most of the perturbations due to the C-terminal 35 amino acid tail were localized in α1, β3, and neighboring regions (**Fig. S7A**). This suggested that the putative α2-helix interacts mostly with the B-fold in these contiguous regions. Mutations R72C and R83C were made at the N- and C-terminal ends of the α2 region in separate samples of $S_{b1}$ and these proteins were derivatized with the stable nitroxide spin label MTSL. Paramagnetic relaxation enhancement (PRE) measurements (**Fig. S7B**) showed significant decreases in amide peak intensity over the α1 and β3 regions for the B-core of $S_{b1}$, consistent with the chemical shift perturbation data. Furthermore, the PRE intensity profiles were similar regardless of which end of the α2 region the spin label resided. This suggests that docking of the α2 region against the B-folded core of $S_{b1}$ is in exchange between multiple states, providing a plausible explanation for why most of the α2 amide resonances are not detectable. Structures for $S_{b1}$ were re-calculated using additional weak (<20Å) PRE restraints, showing an ensemble with a well-defined B-core that has a putative α2-helix packed against it loosely (**Fig. S6D, Table 2**). Steady-state $\{^1H\}$-$^{15}N$ heteronuclear NOE data for $S_{b1}$ were consistent with the structure (**Fig. S6E**). In particular, the C-terminal tail becomes more ordered around the α2 region, although these heteronuclear NOE values (0.4-0.7) are still below those of well-ordered regions (>0.8). Thus, the structure of $S_{b1}$ may be viewed as a transitory state between the S- and B-folds. With two $\alpha$-helices packed against a 4-stranded $\beta$-sheet, $S_{b1}$ has the same overall two-layer $\alpha/\beta$-sandwich architecture as the S-fold but differs in the topological arrangement of secondary structures.

**Structure of** $S_{b2}$**.** The three dimensional structure of $S_{b2}$ contains four $\beta$-strands and two $\alpha$-helices and has the general features of the parent S-fold (**Fig. S9B,C, Table 2**). The ordered regions in the structure are residues 1-9 ($\beta$1), 23-32 ($\alpha$1), 43-48 ($\beta$2), 59-65 ($\beta$3), 71-80 ($\alpha$2), and 86-91 ($\beta$4). While the parent S (PDB 1RIS) and $S_{a1}$ structures are very similar, the $S_{b2}$ structure differs from both in a number of ways despite having the same overall topology. The $\alpha$1-helix in $S_{b2}$ is shorter, comprising 10 amino acids compared with 17 amino acids in $S_{a1}$. Also, the $\beta$2-strand forms 4 amino acids later in the $S_{b2}$ polypeptide chain than in $S_{a1}$. The first residue in the $\beta$2-strand of $S_{b2}$, G43, interacts with E65 in the $\beta$3-strand. This represents a two-residue shift in the register of hydrogen bonding between $\beta$2 and $\beta$3 in $S_{b2}$ compared with $S_{a1}$ (**Fig. S4B**). As a result of these differences, the loops connecting $\beta$1 to $\alpha$1 and $\alpha$1 to $\beta$2 are longer in $S_{b2}$ (13 and 10 residues, respectively) than in $S_{a1}$ (5 and 7 residues, respectively). The remainder of the $S_{b2}$ structure encompassing $\beta$1, $\beta$3, $\alpha$2, and $\beta$4 is very similar to $S_{a1}$. Heteronuclear NOE dynamics data for $S_{b2}$ were consistent with the NMR structure (**Fig. S9D**). In particular, the relatively long $\beta$1-$\alpha$1, $\alpha$1-$\beta$2, and $\beta$2-$\beta$3 loops were found to be the most flexible on the ps-ns timescale.


1 Chen, Y. *et al.* Rules for designing protein fold switches and their implications for the folding code. *bioRxiv*, 2021.2005.2018.444643, doi:10.1101/2021.05.18.444643 (2021).

2 He, Y., Chen, Y., Rozak, D. A., Bryan, P. N. & Orban, J. An artificially evolved albumin binding module facilitates chemical shift epitope mapping of GA domain interactions with phylogenetically diverse albumins. *Protein Sci* **16**, 1490-1494 (2007).

3 Alexander, P., Fahnestock, S., Lee, T., Orban, J. & Bryan, P. Thermodynamic analysis of the folding of the Streptococcal Protein G IgG-binding domains B1 and B2: why small proteins tend to have high denaturation temperatures. *Biochemistry* **31**, 3597-3603 (1992).

4 Haglund, E. *et al.* Trimming down a protein structure to its bare foldons: spatial organization of the cooperative unit. *J Biol Chem* **287**, 2731-2738, doi:10.1074/jbc.M111.312447 (2012).
